# Persistent DNA damage reprograms the cutaneous microenvironment into a stress-responsive immunometabolic pro-pigmentary niche

**DOI:** 10.64898/2026.11.16.767063

**Authors:** Ali Nasrallah, Nazim Mechkouri, Walid Mahfouf, Elodie Muzotte, Sergio Oterino-Sogo, Fatima Naji, Joudi El Mir, Mercedes Quintana, Corinne Buré, Benoît Pinson, Sabrina Lacomme, Etienne Gontier, Stéphane Claverol, Alain Taieb, Fanny Morice-Picard, Léa Dousset, Elodie Darbo, Hamid-Reza Rezvani

**Author notes:** Authors contributed equally (Ali Nasrallah, Nazim Mechkouri). **Corresponding Author** Postal address: INSERM U1312, Bordeaux, F-33000 France, Email address.

## Abstract

Chronic ultraviolet (UV) exposure induces progressive pigmentary remodeling of the skin, yet how persistent DNA damage influences tissue-level responses remains poorly understood. Xeroderma pigmentosum complementation group C (XP-C), characterized by defective global genome nucleotide excision repair, provides a model to investigate these processes. Here, integrating chronic UVB irradiation of *Xpc*-deficient mice with spatial transcriptomics, multi-omics profiling, and functional analyses, we identify a coordinated multicellular response associated with persistent hyperpigmentation. UVB-exposed *Xpc*-deficient skin exhibited increased eumelanin accumulation, altered melanosome organization, enhanced melanosome transfer to keratinocytes, and regional melanocyte expansion. Proteomic, metabolomic, and kinomic analyses revealed coordinated alterations in inflammatory, cytoskeletal, redox, metabolic, and pigmentation-associated programs. Spatial transcriptomics further identified distinct keratinocyte states and inflammatory immune niches enriched in hyperpigmented regions, coinciding with increased melanocyte abundance and DNA damage. These findings suggest that pigmentary remodeling occurs within a spatially organized, stress-responsive immunometabolic tissue environment rather than reflecting melanocyte-intrinsic alterations alone. Analyses of human XP-C skin and XPC-deficient primary human melanocytes revealed overlapping pigmentation-associated, inflammatory, and cytoskeletal alterations, supporting the relevance of these observations to human disease. Together, our findings reveal a previously underappreciated relationship between persistent DNA damage and spatially coordinated multicellular remodeling, providing a framework for understanding how defective genome maintenance is associated with chronic pigmentary changes and altered tissue homeostasis. These findings may also offer insights into the mechanisms underlying common acquired hyperpigmentary disorders.

**Bullet:**

- Persistent DNA damage reprograms the cutaneous microenvironment into a multicellular stress-responsive immunometabolic tissue state.
- Spatial transcriptomics identifies coordinated activation of melanocytes, keratinocytes and immune cells underlying chronic UV-induced hyperpigmentation.
- Complementary multi-omics analyses identify oxidative stress, inflammatory signaling, cytoskeletal remodeling, and metabolic rewiring as key features of persistent hyperpigmentation.
- XPC deficiency provides as an accelerated model of chronic UV-induced pigmentary remodeling, revealing mechanisms potentially relevant to acquired hyperpigmentation.

## Introduction

The skin constitutes the body’s primary barrier against environmental insults and is continuously exposed to solar ultraviolet (UV) radiation ^1^. Among the adaptive mechanisms that protect against chronic photodamage, cutaneous pigmentation is one of the most effective. Following UV exposure, melanocytes increase melanin synthesis and transfer melanosomes to neighboring keratinocytes, thereby limiting DNA damage and oxidative stress within the epidermis ^2,3^. While constitutive pigmentation is largely determined by genetic background, UV-induced pigmentation represents a dynamic adaptive process that integrates DNA damage responses, intercellular communication and tissue remodeling ^4^.

Repeated UV exposure induces a spectrum of pigmentary responses ranging from transient tanning to persistent hyperpigmented lesions such as solar lentigo ^5^. Unlike physiological tanning, which is rapidly induced and largely reversible, chronic UV-induced hyperpigmentation is associated with long-lasting alterations in epidermal architecture, melanocyte distribution, keratinocyte behavior and local inflammatory responses ^6^. Although numerous molecular regulators of melanogenesis have been identified ^7^, the mechanisms by which persistent UV-induced damage coordinates these multicellular adaptations and establishes long-lasting pigmentary remodeling remain poorly understood. An unresolved question is whether persistent DNA damage itself acts as the initiating signal that drives this tissue-wide remodeling. Addressing this question has been challenging because most experimental models rely on acute UV exposure or isolated cell systems ^8^, which fail to reproduce the complex interactions among melanocytes, keratinocytes, immune cells and stromal cells that collectively determine the pigmentation phenotype.

Xeroderma pigmentosum (XP) provides a unique opportunity to investigate this question. XP is a rare inherited disorder caused by defects in nucleotide excision repair (NER), resulting in extreme photosensitivity, early-onset pigmentary abnormalities and a dramatically increased risk of skin cancer ^9,10^. Among XP complementation groups, XP-C arises from mutations in XPC, the primary sensor of helix-distorting DNA lesions during global genome NER ^9^. Consequently, XPC has been studied predominantly for its roles in DNA damage recognition, genome stability and photocarcinogenesis ^11–13^. However, one of the earliest and most characteristic clinical manifestations of XP-C is the development of relatively stable hypo- and hyperpigmented macules in chronically sun-exposed skin, often appearing during early childhood ^14^. These pigmentary abnormalities share several clinical and histopathological features with common UV-associated pigmentary disorders ^14^. Notably, some XP-C patients develop lesions histologically resembling solar lentigines ^14,15^, suggesting that XP-C may represent an accelerated model of chronic UV-induced pigmentary remodeling. Importantly, pigmentary abnormalities in patients with XP-C can remain remarkably stable over several years ^14^ (see also **Fig. 1A**), including under stringent photoprotection. Such persistence raises the possibility that defective repair of UV-induced DNA damage establishes durable alterations in the cutaneous tissue environment rather than merely eliciting a transient melanogenic response. Whether these persistent changes arise from melanocyte-intrinsic alterations or from broader tissue remodeling involving sustained inflammatory signaling, metabolic adaptation, altered cellular states, and disrupted intercellular communication remains unclear.

**Figure 1.**
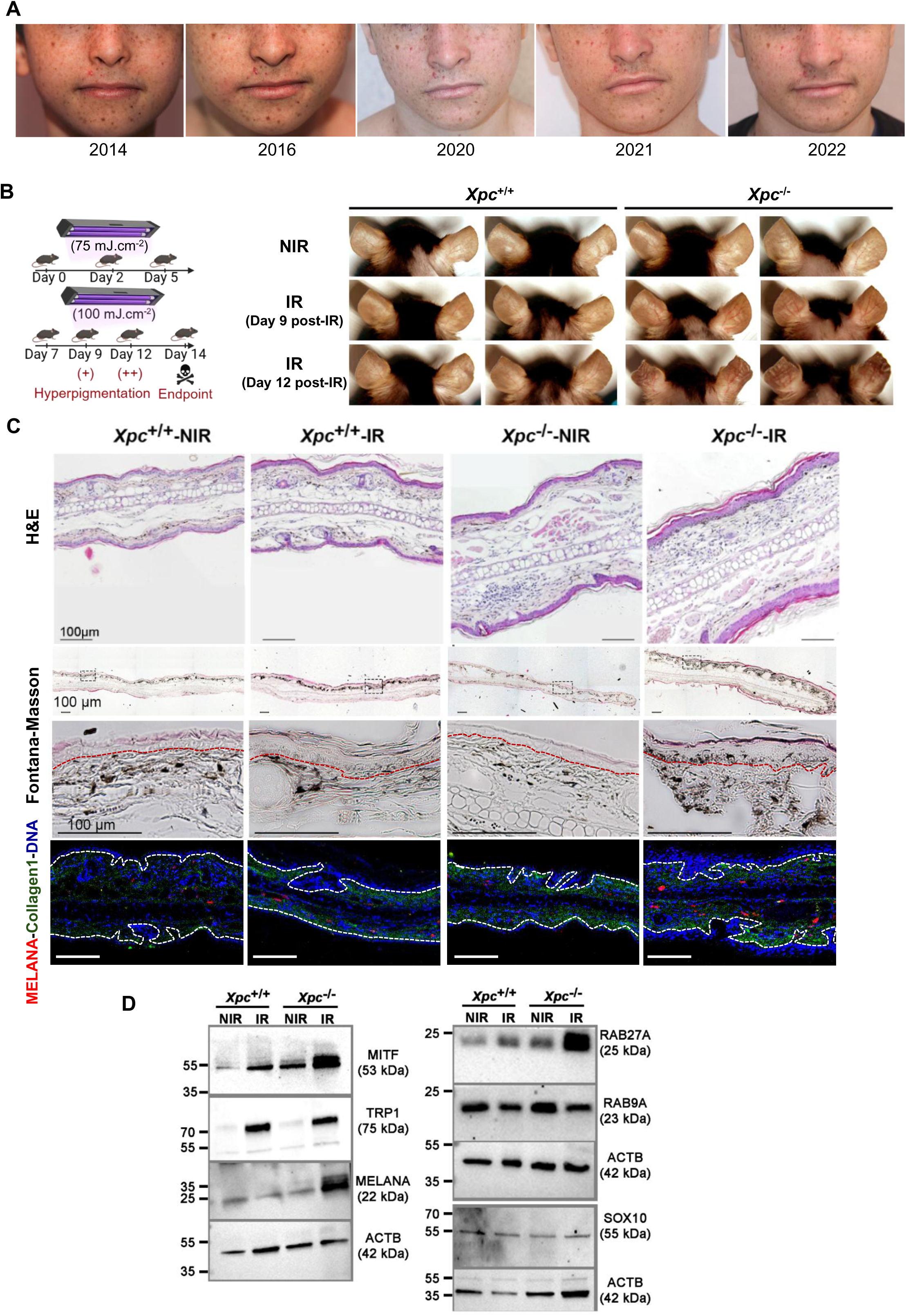
XPC deficiency drives rapid UVB-induced hyperpigmentation through melanocyte redistribution and activation of melanogenic programs. (**A**) Representative clinical photographs of an XP-C patient showing persistent hyper- and hypopigmented macules on sun-exposed skin over an extended follow-up period. The spatial distribution of pigmentary lesions remained largely stable over time. Written informed consent for publication was obtained from the patient or legal guardians. The study was approved by the local ethics committee of the University Hospital of Bordeaux, France (CE-GR-2017/006). (**B**) Experimental design of the chronic UVB irradiation model (left). Wild-type (*Xpc*^+/+^) and Xpc-deficient (*Xpc*^−/−^) mice were exposed to repeated UVB irradiation (IR; 75 mJ/cm^2^ during week 1 and 100 mJ/cm^2^ during week 2) or maintained under non-irradiated conditions (NIR). Representative macroscopic images (right) illustrate the progressive development of ear hyperpigmentation following irradiation. (**C**) Histological characterization of ear skin from *Xpc*^+/+^ and *Xpc*^−/−^ mice under NIR and IR conditions. Hematoxylin and eosin (H&E) staining reveals epidermal architecture and UVB-induced epidermal thickening (top panels). Fontana–Masson staining highlights melanin deposition and its distribution within the skin. Higher-magnification images of the boxed regions are shown below. Imaging mass cytometry (Hyperion) using antibodies against Melan-A (red) and collagen I (green), together with a nuclear counterstain (blue), reveals the distribution of melanocytes within the epidermis and dermis (bottom panels). Red dashed lines in the Fontana–Masson images and white dashed lines in the immunofluorescence images indicate the dermal– epidermal junction. Scale bars, 100 μm. (**D**) Representative immunoblot analysis of melanogenic and melanosome-associated proteins in ear lysates from *Xpc*^+/+^ and *Xpc*^−/−^ mice. β-actin was used as a loading control. Molecular weights (kDa) are indicated. NIR, nonirradiated; IR, UVB-irradiated.

Here, we establish a chronic UVB irradiation model using Xpc-deficient mice that reproduces the hyperpigmented phenotype observed in XP-C patients. By integrating histology, ultrastructural imaging, quantitative melanin analyses, functional melanosome-transfer assays, quantitative proteomics, metabolomics, kinome profiling, and Xenium spatial transcriptomics, we investigate the multicellular mechanisms linking persistent DNA damage to pigmentary remodeling. We further validate key features of this phenotype in skin biopsies from XP-C patients and in CRISPR/Cas9-engineered human melanocytes, supporting the translational relevance of the mechanisms identified in the mouse model.

Our findings demonstrate that XPC deficiency does not simply enhance melanocyte-autonomous melanogenesis. Instead, persistent DNA damage drives coordinated reprogramming of melanocytes, keratinocytes and immune cells, leading to cytoskeletal remodeling, inflammatory activation, metabolic adaptation, enhanced melanosome transfer and tissue-wide hyperpigmentation. These results identify persistent DNA damage as a central regulator of multicellular skin adaptation to chronic UV exposure and establish a mechanistic framework that may extend beyond XP-C to common acquired hyperpigmentary disorders.

## Results

### XPC deficiency drives rapid UVB-induced hyperpigmentation through melanocyte redistribution and altered melanosome dynamics

Alteration in the skin pigmentary phenotype associated with the effects of UV radiation is one of the first clinical manifestations in most of NER disorders. Our clinical evolution of pigmentation in affected XP-C individuals revealed that hyper- and hypopigmented macules appeared early in life, typically before two years of age, and remained remarkably stable in their spatial distribution over prolonged follow-up (**Fig. 1A**). Although these lesions share certain clinical features with common UV-associated pigmentary disorders, including solar lentigines and idiopathic guttate hypomelanosis^14^, the mechanisms driving their formation remain poorly understood due to the lack of an appropriate model. To model XP-associated pigmentary abnormalities, XPC-deficient mice were subjected to a short-term chronic UVB irradiation protocol (**Fig. 1B**). Under non-irradiated (NIR) conditions, *Xpc*^−/−^ mice were indistinguishable from wild-type littermates and showed no overt pigmentary abnormalities. In contrast, repeated UVB exposure rapidly induced visible hyperpigmentation in *Xpc*^−/−^ ears, becoming apparent by day 9 and markedly pronounced by day 12, whereas only minimal pigmentation changes were observed in irradiated wild-type controls (**Fig. 1B**).

Hematoxylin and eosin staining revealed epidermal thickening following UVB exposure, whereas Fontana–Masson staining demonstrated markedly increased melanin deposition in irradiated *Xpc*^−/−^ ears compared with the other experimental groups (**Fig. 1C**). Notably, under nonirradiated conditions, melanin deposits and Melan-A-positive melanocytes were predominantly localized within the dermal compartment, irrespective of genotype. Following UVB irradiation, both melanin deposits and Melan-A-positive cells were observed in the dermal and epidermal compartments, with a more pronounced accumulation in *Xpc*^−/−^ mice (**Fig. 1C**).

To determine whether these morphological changes were associated with activation of melanogenic programs, we examined the expression of key pigmentation regulators. Immunoblot analyses revealed robust induction of MITF, TYRP1 and MELAN-A in irradiated *Xpc*^−/−^ skin compared with all other experimental groups (**Fig. 1D**). Increased expression of RAB27A further suggested enhanced melanosome trafficking and potentially enhanced melanosome transfer to keratinocytes (**Fig. 1D**).

Ultrastructural analyses provided additional evidence for profound alterations in pigment organelle biology. Transmission electron microscopy revealed a marked accumulation of melanosomes throughout *Xpc*^−/−^ epidermis, together with the appearance of abundant multimelanosome vesicles following UVB exposure (**Fig. 2A–D**). These structures were detected in dermal melanocytes, epidermal melanocytes, keratinocytes and even within the cornified layer, supporting extensive redistribution and transfer of melanin throughout the epidermis (**Fig. 2D**).

**Figure 2.**
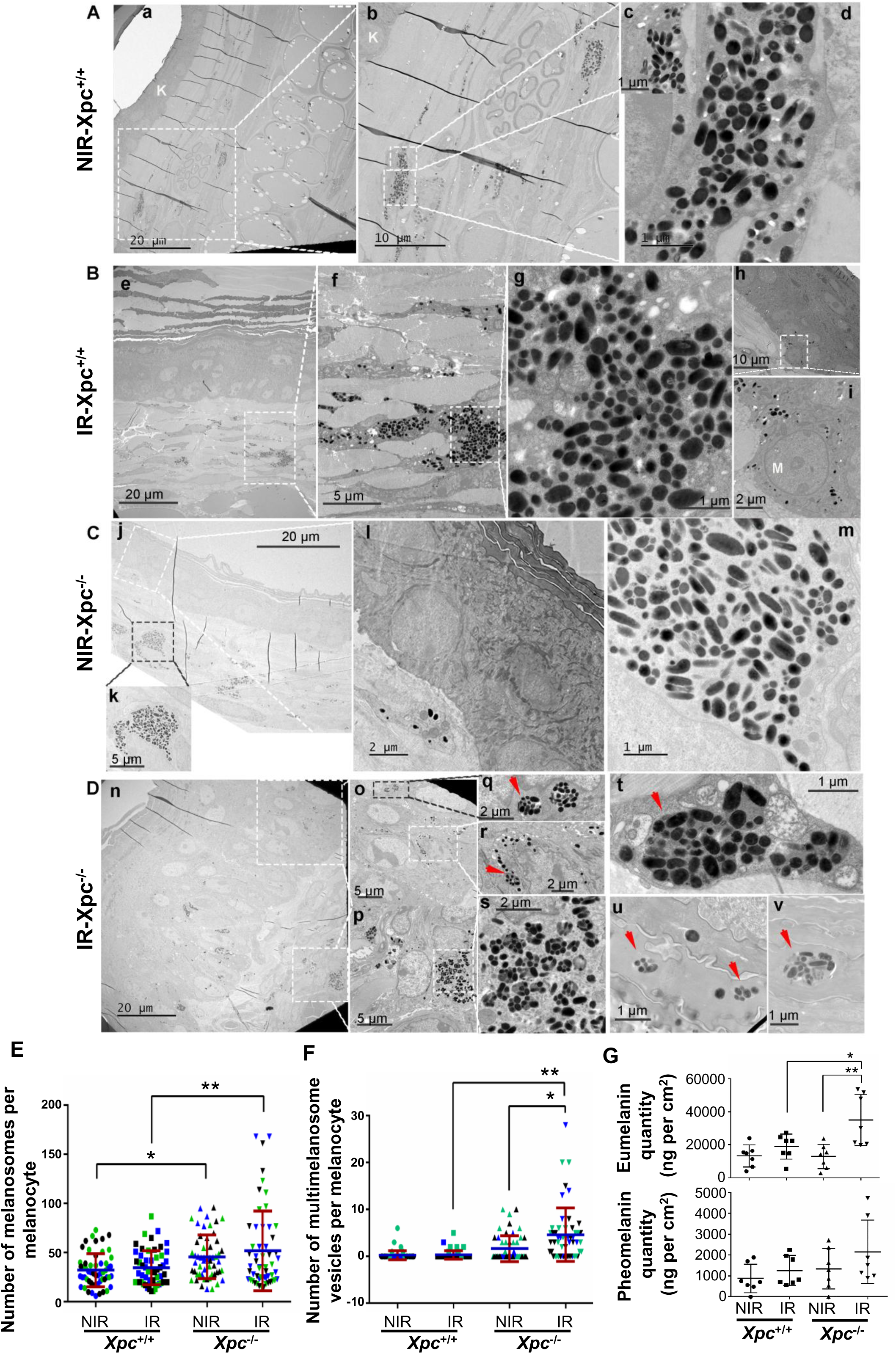
Electron microscopy analyses revealed abnormal melanosome organization and enhanced pigment transfer in UVB-irradiated *Xpc*^−/−^ skin. (**A-D**) Representative transmission electron microscopy (TEM) images of ear skin from different conditions. (**A**) TEM images of non-irradiated *Xpc*^+/+^ ear skin showing dermal melanocytes containing predominantly dispersed mature melanosomes. Insets (b–d) illustrate melanosome morphology and intracellular localization at higher magnification. K, keratinocyte. (**B**) TEM images of irradiated *Xpc*^+/+^ ear skin showing focal melanosome clustering within dermal melanocytes (e–g) and occasional melanocytes and melanosomes within the epidermis (h–j). M, melanocyte. (**C**) Representative TEM images of non-irradiated *Xpc*^−/−^ ear skin demonstrating dermal melanocytes and melanosome distribution patterns (j–m). (**D**) TEM analysis of irradiated *Xpc*^−/−^ ear skin showing extensive accumulation of melanosomes and multimelanosome vesicles (membrane-bound pigment globules containing multiple melanosomes) within epidermal melanocytes and keratinocytes (n, o, q, r), dermal melanocytes (p, s, t), and the cornified layer (u, v). Scale bars are indicated in each panel. (**E-F**) Quantification of total number of melanosomes (**E**) and multimelanosome-containnig vesicles (**F**) per melanocyte in *Xpc*^+/+^ and *Xpc*^−/−^ mice under NIR and IR conditions. Melanosomes and multimelanosome-containing vesicles were quantified in 55 melanocytes per experimental condition, randomly selected from tissue sections obtained from three independent mice per group and examined by transmission electron microscopy. Individual melanocyte measurements from each mouse are shown in a different color (blue, green, or black). Blue lines indicate the mean, and red lines represent the standard deviation (SD) of the 55 melanocyte measurements per condition (n = 3 biologically independent mice per group). (**G**) Quantification of eumelanin (top) and pheomelanin (bottom) content in ear skin samples. Eumelanin levels were significantly elevated in irradiated *Xpc*^−/−^ mice, whereas pheomelanin content did not differ significantly among experimental groups. Data are presented as mean ± SD (n = 7 biologically independent mice per group). Statistical significance in E-G was determined using two-way ANOVA on mouse-level data, followed by Šídák’s multiple comparisons test. *P < 0.05; **P < 0.01. NIR, nonirradiated; IR, UVB-irradiated.

Quantitative analyses revealed a significant increase in the total number of melanosomes per melanocyte in *Xpc*^−/−^ mice under both nonirradiated and UVB-irradiated conditions compared with their respective wild-type controls (**Fig. 2E**), suggesting that XPC deficiency promotes melanosome accumulation independently of UVB exposure. In contrast, multimelanosome-containing vesicles were significantly more abundant in irradiated *Xpc*^−/−^ melanocytes than in either nonirradiated *Xpc*^−/−^ or irradiated wild-type melanocytes (**Fig. 2F**). These findings suggest that, beyond increasing melanosome abundance, XPC deficiency is associated with altered melanosome organization, particularly following UVB irradiation.

Given the distinct photobiological properties of eumelanin and pheomelanin, we next quantified their abundance. UVB irradiation induced a significantly greater increase in eumelanin content in *Xpc*^−/−^ ears than in wild-type controls (**Fig. 2G**). Indeed, eumelanin levels were comparable between nonirradiated genotypes but significantly elevated in irradiated *Xpc*^−/−^ mice compared with both nonirradiated *Xpc*^−/−^ and irradiated wild-type mice. In contrast, pheomelanin levels remained largely unchanged across experimental groups, supporting a preferential enhancement of eumelanin accumulation in UVB-irradiated XPC-deficient skin.

To directly assess whether XPC deficiency influences melanosome transfer to keratinocytes, primary melanocytes were isolated from non-irradiated *Xpc*^+/+^ and *Xpc*^−/−^ mouse ears (**Fig. S1A**) and then labeled with the melanosome-specific fluorescent reporter MelanoTag ^16^. These red fluorescent melanocytes were subsequently co-cultured with GFP-expressing mouse keratinocytes and then exposed twice to UVB irradiation. Time-lapse imaging visualized the transfer of MelanoTag-positive melanosomes from UVB-irradiated *Xpc*^−/−^ melanocytes to neighboring keratinocytes (**Fig. S1B**). Five hours after the second irradiation, limited melanosome transfer was observed in wild-type co-cultures, whereas co-cultures containing *Xpc*^−/−^ melanocytes exhibited markedly increased accumulation of MelanoTag-positive vesicles within keratinocytes (**Fig. S1C**), suggesting that XPC deficiency potentiates UVB-induced melanosome transfer.

Collectively, these findings demonstrate that XPC deficiency profoundly alters the cutaneous pigmentary response to UVB, promoting melanocyte redistribution, enhanced melanogenic activity, altered melanosome biogenesis and organization, increased melanosome transfer, and selective accumulation of photoprotective eumelanin.

### Multi-omics profiling identifies convergent activation of melanogenic, vesicle trafficking, inflammatory and metabolic rewiring in UVB-irradiated Xpc-deficient skin

To investigate the molecular mechanisms underlying the exaggerated pigmentation phenotype observed in UVB-irradiated Xpc-deficient skin, we performed an integrated multi-omics analysis combining quantitative proteomics, metabolomics, and functional kinome profiling.

Label-free quantitative proteomics of ear skin samples identified 4,163 proteins across all experimental groups (n = 5 mice per group). Principal component analysis demonstrated a clear segregation of irradiated *Xpc*^−/−^ samples from all other conditions, indicating extensive proteome remodeling driven by the combined effects of XPC deficiency and chronic UVB exposure (**Fig. 3A**). Consistent with this observation, 809 and 761 differentially expressed proteins (DEPs) were identified when irradiated *Xpc*^−/−^ skin was compared with non-irradiated *Xpc*^−/−^ skin and irradiated *Xpc*^+/+^ skin. In contrast, only 22 proteins differed between non-irradiated *Xpc*^−/−^ and *Xpc*^+/+^ mice, whereas 151 proteins were altered following irradiation of wild-type mice (**Fig. S2A**), indicating that UVB exposure unmasks profound molecular consequences of XPC deficiency.

**Figure 3.**
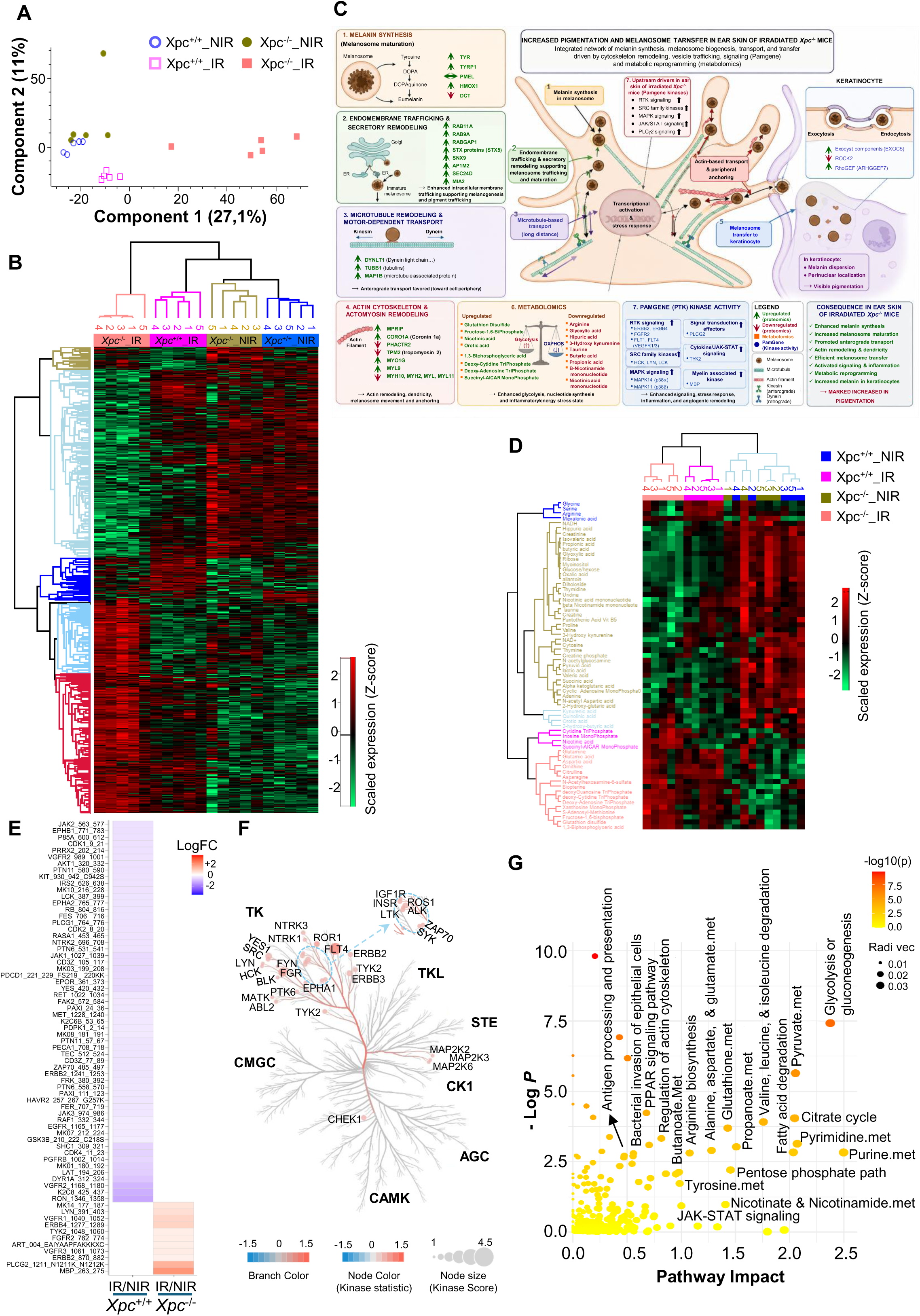
Integrated proteomic, metabolomic and kinome profiling identifies convergent pathways driving hyperpigmentation in XPC-deficient skin. (**A**) Principal component analysis (PCA) of label-free quantitative proteomic profiles obtained from ear skin samples of non-irradiated (NIR) and irradiated (IR) *Xpc*^+/+^ and *Xpc*^−/−^ mice (n = 5 per group). (**B**) Unsupervised hierarchical clustering heatmap of differentially abundant proteins (rows) across all experimental groups (columns). The green-to-red color scale represents relative protein abundance following normalization and Z-score transformation, with green indicating lower abundance and red indicating higher abundance. Hierarchical clustering was performed using Spearman correlation and complete linkage in Perseus. Biological replicates clustered according to genotype and irradiation status, revealing distinct group-specific proteomic profiles. (**C**) Integrated mechanistic model summarizing major molecular alterations identified by proteomics, metabolomics and kinome profiling in irradiated *Xpc*^−/−^ ear skin. Differentially regulated proteins, metabolites and kinase pathways are organized according to their predicted roles in melanogenesis, melanosome biogenesis and trafficking, cytoskeletal remodeling, inflammation and metabolic adaptation. Red and green indicate upregulated and downregulated molecules, respectively. (**D**) Semi-targeted metabolomic profiling of ear skin samples (n = 5 per group). Heatmap showing relative abundance of significantly altered metabolites (rows) across experimental groups (columns). Hierarchical clustering was performed using Spearman correlation and complete linkage. The green-to-red color scale represents relative metabolite abundance after normalization and Z-score transformation. (**E**) Functional protein tyrosine kinase (PTK) profiling using PamGene peptide arrays. Differential peptide phosphorylation was determined by comparing irradiated (IR) and non-irradiated (NIR) ear skin from *Xpc*^+/+^ and *Xpc*^−/−^ mice. Heatmap colors represent log fold-change (LogFC) values for significantly altered phosphopeptides, revealing distinct phosphorylation signatures induced by UVB irradiation in wild-type and XPC-deficient skin. (**F**) Kinome tree representation of predicted upstream kinase activity generated by Upstream Kinase Analysis (UKA) of PamGene PTK datasets and visualized using CORAL. Branch color represents the predicted direction of kinase activity, node color corresponds to the kinase statistic, and node size reflects the kinase score. (**G**) Integrated pathway analysis of proteomic, metabolomic and kinome datasets using the Joint Pathway Analysis module of MetaboAnalyst. Differentially expressed proteins, metabolites and predicted upstream kinases were jointly mapped onto KEGG pathways and analyzed using topology-based pathway enrichment. Each circle represents an enriched biological pathway. The x-axis indicates pathway impact, whereas the y-axis represents pathway enrichment significance (−log10 *P* value). Circle color reflects statistical significance and circle size corresponds to pathway impact. NIR, nonirradiated; IR, UVB-irradiated.

Unsupervised hierarchical clustering robustly separated all samples according to genotype and irradiation status, with complete segregation of biological replicates within each experimental group (**Fig. 3B**). Functional enrichment analyses using Ingenuity Pathway Analysis (IPA) (www.ingenuity.com) identified immune cell trafficking, skin development and function, and tissue morphology among the most significantly altered biological processes in irradiated *Xpc*^−/−^ skin (**Fig. S2B**). Consistently, disease and function analyses highlighted inflammatory responses, metabolic disorders, dermatological conditions, and cancer-related pathways as major molecular signatures associated with the hyperpigmented phenotype (**Fig. S2C**). Pathway analyses further revealed extensive activation of inflammatory and interferon-associated signaling networks, extracellular matrix remodeling, vesicle-mediated transport, and cellular redox pathways (**Fig. S2D**). Concomitantly, glycolysis and nucleotide metabolism were predicted to be activated, whereas oxidative phosphorylation and fatty acid β-oxidation were suppressed, indicating a profound metabolic shift toward stress-adapted energy utilization (**Fig. S2E**). Notably, several proteins associated with pigmentation-related processes were differentially abundant, including proteins involved in melanin synthesis (TYR, TYRP1, and HMOX1), potential regulators of melanosome transfer (EXOC6 and ARHGEF7), and components of endomembrane trafficking and secretory pathways (RAB11A, RAB9A, RABGAP1, STX5, SNX9, AP1M2, SEC24D, and MIA2). Additional alterations involved proteins associated with microtubule organization and motor-dependent transport (DYNLLT1, TUBB1, and MAP1B), as well as actin cytoskeleton and actomyosin remodeling. Within the latter group, MPRIP, CORO1A, MYO1G, and MYL9 were increased, whereas PHACTR2, TPM2, MYH10, MYH2, MYL1, and MYL11 were decreased (**Fig. 3C**). Together, these alterations suggest coordinated enhancement of melanin synthesis, melanosome maturation, intracellular trafficking and pigment transfer.

To examine the metabolic consequences of XPC deficiency and chronic UVB exposure, we next performed semi-targeted metabolomic profiling. Among 139 detected metabolites, unsupervised analyses revealed clear separation of irradiated and non-irradiated groups, with irradiated *Xpc*^−/−^ samples displaying the most distinct metabolic profile (**Fig. 3D**). Pathway enrichment analyses identified significant perturbations in purine and pyrimidine metabolism, glycolysis, nicotinate and nicotinamide metabolism, amino acid metabolism, and the metabolism of taurine, hypotaurine, butanoate and propanoate (**Fig. 3D** and **Fig. S3A**). Consistent with the proteomic analyses, irradiated *Xpc*^−/−^ skin exhibited accumulation of glycolytic intermediates, including fructose-1,6-bisphosphate and 1,3-bisphosphoglycerate, together with increased levels of deoxyribonucleotide triphosphates (dNTPs), supporting activation of glycolysis and nucleotide metabolism (**Fig. 3D and Fig. S3B**). Of particular interest, marked reductions in taurine, butanoate, propanoate and kynurenine-related metabolites, coupled with increased glutathione disulfide levels, indicated disruption of redox homeostasis and persistence of oxidative stress signaling in irradiated *Xpc*^−/−^ skin (**Fig. 3D and Fig. S3B**), collectively suggesting disruption of redox homeostasis and persistence of oxidative stress signaling.

Canonical pathway analysis of differential metabolites using IPA predicted activation of nucleotide biosynthesis, MAP kinase signaling, and immune-associated kinase networks, including FLT3-related pathways, accompanied by extensive remodeling of redox balance and cellular energy metabolism (**Fig. S3C**). Such metabolic alterations are consistent with a pro-inflammatory microenvironment and sustained melanogenic activation.

To investigate upstream signaling events associated with these molecular alterations, we performed functional protein tyrosine kinase (PTK) profiling using PamGene arrays. Comparison of irradiated and non-irradiated skin revealed distinct phosphorylation signatures between genotypes. Whereas UVB irradiation predominantly reduced peptide phosphorylation in wild-type skin, *Xpc*^−/−^ skin exhibited a marked increase in phosphorylation of multiple PTK substrates, indicating activation of a distinct kinase signaling program (**Fig. 3E**). Upstream Kinase Analysis (UKA) further predicted coordinated activation of several receptor tyrosine kinase and non-receptor tyrosine kinase families, such as ERBB2, FLT4, IGF1R, ALK, FGR, and NTRK1 (**Fig. 3F**). In parallel, activation of the MAP2K2–MAP2K3–MAP2K6 signaling module, which lies upstream of p38 MAPK (MAPK14), was predicted, suggesting engagement of stress-responsive MAP kinase signaling (**Fig. 3F**). Kinome tree visualization demonstrated coordinated activation of both receptor and intracellular tyrosine kinase families, together with STE kinases, highlighting coordinated activation of signaling networks governing growth factor signaling, inflammatory responses, cytoskeletal remodeling, vesicle trafficking and melanocyte biology (**Fig. 3C, F**).

To obtain a systems-level view of these molecular alterations, proteomic, metabolomic and kinome datasets were integrated using the Joint Pathway Analysis module of MetaboAnalyst. This analysis identified coordinated perturbations in multiple interconnected metabolic and signaling pathways, with glycolysis/gluconeogenesis, pyruvate metabolism, purine and pyrimidine metabolism, the tricarboxylic acid cycle, fatty acid degradation, and the pentose phosphate pathway emerging as the most significantly impacted networks (**Fig. 3G**). Additional alterations involved amino acid metabolism, nicotinate/nicotinamide metabolism, regulation of the actin cytoskeleton, and JAK– STAT signaling. Together, glutathione metabolism and the pentose phosphate pathway emerged as major nodes of pathway convergence, further supporting a central role for oxidative stress adaptation and redox remodeling in the development of the hyperpigmented phenotype.

Collectively, integration of the proteomic, metabolomic, and kinomic datasets revealed convergence of molecular alterations across pathways associated with melanocyte activation, vesicle trafficking, cytoskeletal remodeling, inflammatory signaling, and oxidative stress responses (**Fig. 3C**). These findings suggest that the enhanced melanogenic phenotype associated with persistent DNA damage in XPC-deficient skin occurs within a broader UVB-induced inflammatory and metabolic tissue response, rather than representing an isolated alteration in melanocyte pigmentation.

### Spatial transcriptomic profiling reveals extensive cellular remodeling in UVB-irradiated *Xpc*^−/−^ skin

To resolve the cell-type-specific alterations underlying the molecular programs identified by bulk multi-omics analyses, we performed Xenium spatial transcriptomic profiling of mouse ear skin across the four experimental conditions, following the analytical workflow outlined in **Fig. 4A**. Following quality control and cell segmentation, samples from 6 nonirradiated *Xpc*^+/+^, 8 irradiated *Xpc*^+/+^, 6 nonirradiated *Xpc*^−/−^, and 7 irradiated *Xpc*^−/−^ mice were retained for downstream analyses. Following normalization, dimensionality reduction, and data integration, pooled single-cell transcriptomic profiles were subjected to unsupervised Leiden clustering. Clustering at resolution 1.1 identified 20 transcriptionally distinct clusters, with silhouette scores (>0.3) supporting the separation of major cellular populations (**Fig. 4B**). Manual annotation based on canonical cell-type-specific marker genes resolved the major skin cell populations, including keratinocytes, melanocytes, immune cells, endothelial cells, fibroblasts, adipocytes, and smooth muscle cells (**Fig. 4C,D**).

**Figure 4.**
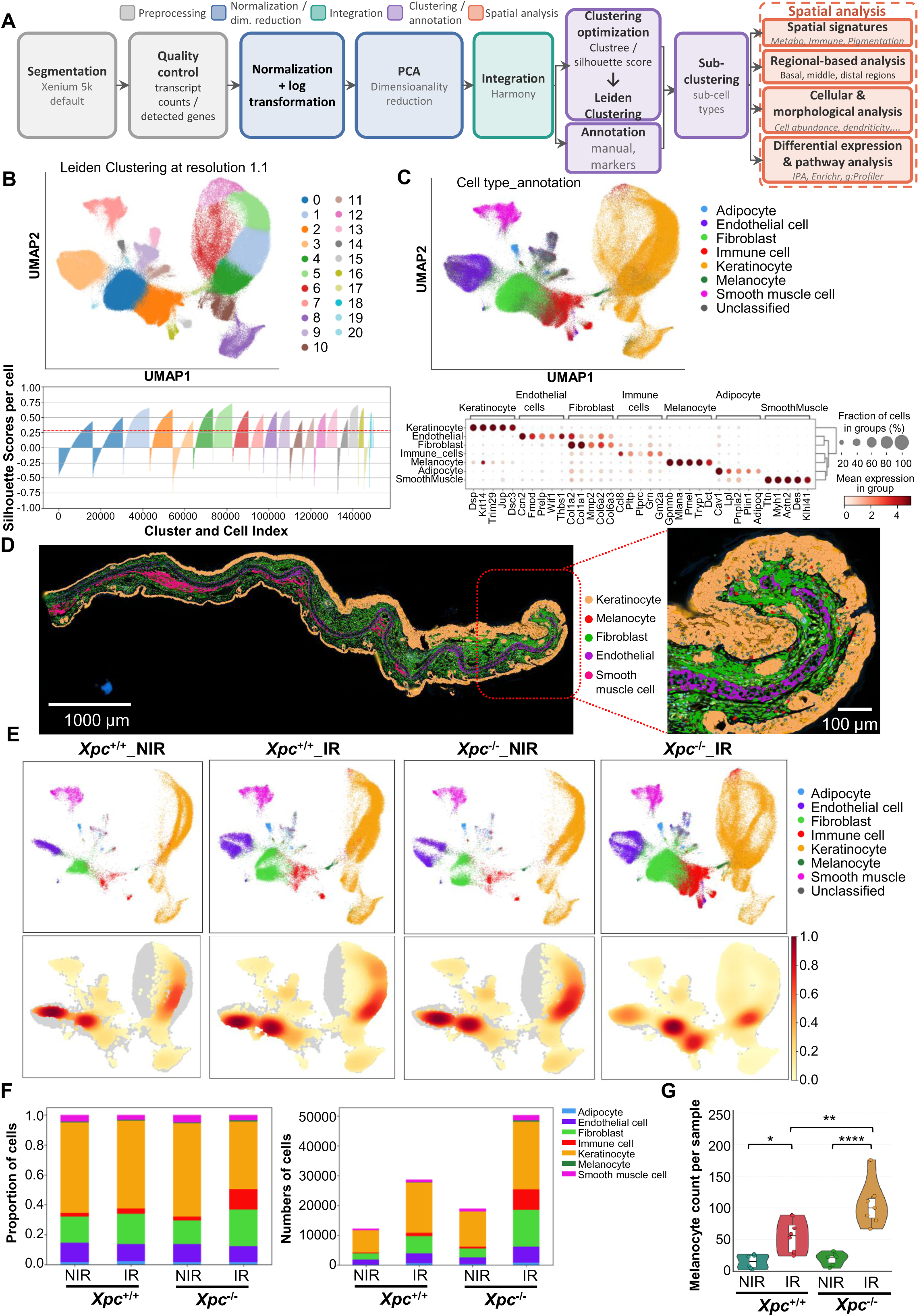
Spatial transcriptomics reveals cellular remodeling of UVB-irradiated Xpc-deficient skin. (**A**) Schematic representation of the analysis pipeline developed for the Xenium 5K dataset generated from the XPC mouse model. The workflow includes preprocessing steps, namely normalization, log-transformation, principal component analysis (PCA), neighbourhood graph construction, Leiden clustering, and UMAP dimensionality reduction, followed by downstream analyses including cell type annotation, visualization, signature heatmaps, spatial mapping, differential expression analysis, niche clustering, and spatially resolved characterization of niche-associated cell types. **(B)** Top panel is UMAP representation of the integrated dataset including all samples from the four experimental conditions, generated using Leiden clustering at a resolution of 1.1. Bottom panel shows silhouette score associated with that clustering. The dashed red line indicates the median silhouette score threshold. **(C)** Top panel is UMAP visualization of annotated cell types identified from canonical marker gene expression. Bottom panel is a dot plot showing expression of representative canonical marker genes across annotated cell types. Dot size indicates the fraction of cells expressing each gene, and color intensity indicates mean expression level. **(D)** Spatial distribution of major cell populations within an irradiated *Xpc*^−/−^ ear section. Keratinocytes (orange), melanocytes (red), fibroblasts (green), and endothelial cells (purple), identified by Xenium-based cell-type annotation, are projected onto their original tissue coordinates. The enlarged view highlights their spatial organization within the distal ear region **(E)** Split UMAP projections showing the distribution of annotated cell types across the four experimental conditions (top), with corresponding density maps illustrating the distribution of cells from each condition within the global UMAP space (bottom). **(F)** Bar plots showing the relative proportions (left) and absolute counts (right) of annotated cell types across experimental conditions. **(G)** Violin plots showing raw melanocyte counts across experimental conditions. Statistical significance was assessed using two-way ANOVA, with *P* < 0.05 considered statistically significant. *P < 0.05; **P < 0.01, *** P < 0.001, ****P < 0.0001. NIR, nonirradiated; IR, UVB-irradiated.

Projection of annotated cells across the four experimental conditions revealed marked remodeling of the cellular composition in UVB-irradiated *Xpc*^−/−^ skin (**Fig. 4E**). Although the global embedding showed partial overlap among the major cell populations, irradiated *Xpc*^−/−^ ears showed expansion of the keratinocyte, melanocyte, and immune-cell compartments (**Fig. 4E**). Notably, subsets of keratinocytes and immune cells from irradiated *Xpc*^−/−^ skin occupied distinct regions of the UMAP space that were not prominent in the other conditions, suggesting condition-associated transcriptional remodeling of these compartments following UVB exposure. Quantification of both relative proportions and absolute cell numbers confirmed a pronounced increase in immune cells, keratinocytes, and melanocytes in irradiated *Xpc*^−/−^ skin compared with the other conditions (**Fig. 4F,G**).

These data demonstrate that UVB exposure in the absence of XPC induces extensive cellular remodeling of the skin microenvironment and provide a spatial framework to investigate how melanocytes, keratinocytes and immune cells contribute to the hyperpigmented phenotype.

### Melanocytes undergo spatial redistribution and transcriptional reprogramming in UVB-irradiated *Xpc*-deficient skin

Having established the global cellular composition of the skin, we next focused on melanocytes to determine how XPC deficiency affects their spatial distribution and phenotype following chronic UVB exposure. High-resolution multimodal Xenium imaging combined with spatial mapping of melanocyte-associated transcripts (*Pmel*, *Mlana*, and *Dct*) identified melanocytes within the dermal compartment of non-irradiated *Xpc*^+/+^ and *Xpc*^−/−^ ears (**Fig. S4A**). Following UVB irradiation, melanocytes were additionally detected within the epidermal compartment, with this redistribution being particularly prominent in *Xpc*^−/−^ skin (**Fig. S4A and Fig. 5A**). This spatial redistribution was consistent with our ultrastructural observations (see **Fig. 2A**) and paralleled the increased epidermal melanin deposition detected by Fontana–Masson staining (see **Fig. 1C**). High-resolution transcript mapping further revealed melanocyte-associated transcripts distributed along elongated dendritic structures, including projections that were not readily delineated by membrane staining alone (**Fig. 5A**). Consistent with this morphological remodeling, the proportion of dendritic melanocytes was increased in irradiated *Xpc*^−/−^ skin compared with the other experimental groups (**Fig. S4B**). Irradiated *Xpc*^−/−^ melanocytes also displayed increased cell area, together with higher transcript counts and a greater number of detected genes per cell compared with non-irradiated *Xpc*^−/−^ melanocytes (**Fig. S4C, D**).

**Figure 5.**
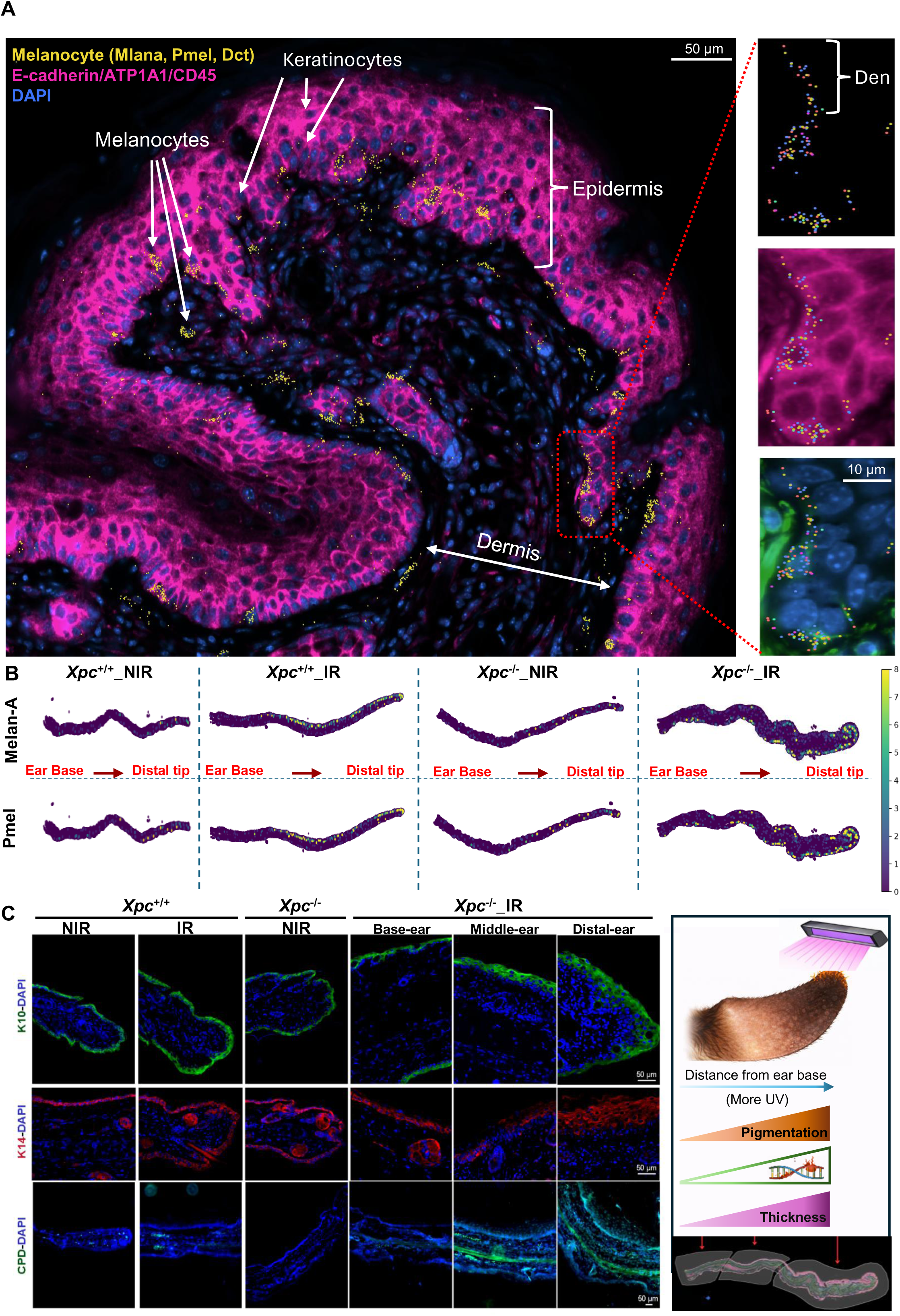
Spatial transcriptomics reveals melanocyte redistribution and a proximal-to-distal pigmentation gradient in UVB-irradiated *Xpc*-deficient ears. (**A**) Representative high-resolution Xenium multimodal image of the distal region of an irradiated *Xpc*^−/−^ ear. DAPI (blue) and E-cadherin/ATP1A1/CD45 (magenta) are shown together with melanocyte-associated transcripts (*Pmel*, *Mlana*, and *Dct*; yellow puncta), illustrating the distribution of melanocytes within the epidermal and dermal compartments. Enlarged views (right) highlight individual melanocytes and their dendritic projections. Melanocyte-associated transcripts are displayed as *Dct* (yellow), *Mlana* (red), *Pmel* (blue), *Tyrp1* (violet), and *Mitf* (pink), overlaid with E-cadherin/ATP1A1/CD45 (magenta), αSMA/vimentin (green), or DAPI (blue), as indicated. Ker, keratinocytes; Mel, melanocytes; Ep, epidermis; Der, dermis; Den, dendrite. **(B)** Representative Xenium spatial maps showing the distribution of melanocyte-associated *Mlana* and *Pmel* transcripts along the longitudinal axis of ears from non-irradiated (NIR) and irradiated (IR) *Xpc*^+/+^ and *Xpc*^−/−^ mice. The arrow indicates the longitudinal orientation of the ear, from the basal region toward the distal tip. **(C)** Immunofluorescence analysis illustrating the spatial gradient of UV-induced tissue remodeling along the longitudinal ear axis. Ear cross-sections from non-irradiated and irradiated *Xpc*^+/+^ and *Xpc*^−/−^ mice were stained for keratinocyte differentiation marker K10 (green), basal keratinocyte marker K14 (red), cyclobutane pyrimidine dimers (CPDs, green) and DAPI (blue). The schematic summarizes the experimental geometry, illustrating the progressive increase in effective UVB exposure from the ear base toward the distal tip, which correlates with increased CPD accumulation, epidermal thickening, and pigmentation. NIR, nonirradiated; IR, UVB-irradiated.

Spatial analysis further revealed increased expression of melanocyte-associated markers, including *Mlana* and *Pmel*, together with higher melanin synthesis and melanosome biogenesis signature scores following irradiation, with the strongest changes observed in *Xpc*^−/−^ skin (**Fig. 5B and Fig. S5A,B**). Notably, these pigmentation-associated signals showed a spatial gradient along the irradiated ear, increasing from the basal region toward the distal tip (**Fig. 5B and Fig. S5A,B**). This gradient paralleled progressive increases in epidermal thickness, accumulation of UVB-induced cyclobutane pyrimidine dimers (CPDs), and visible pigmentation along the longitudinal ear axis (**Fig. 5C**). Because the distal portion of the ear was positioned closer to the UVB source during irradiation, these findings indicate that the experimental configuration generated a longitudinal gradient of effective UVB exposure and photodamage, providing an internal spatial model in which increasing levels of DNA damage are associated with progressive epidermal remodeling and pigmentation (**Fig. 5C and Fig. S5A–C**). Accordingly, subsequent spatial analyses were performed separately in the basal, middle, and distal ear regions, whose dimensions were comparable across experimental groups (**Fig. S5C**). Interestingly, keratinocytes from the basal and distal regions were predominantly assigned to distinct transcriptional subpopulations, whereas both populations were represented within the middle region, further supporting spatial heterogeneity in keratinocyte states along the ear axis (**Fig. S5D**).

Regional quantification confirmed a marked increase in melanocyte abundance in irradiated *Xpc*^−/−^ ears, with the greatest increase observed in the distal ear region (**Fig. 6A**). A more modest increase was also observed in the distal ear region of irradiated wild-type skin. Similar results were obtained after normalizing melanocyte abundance to keratinocyte number, although the magnitude of the difference was attenuated, indicating that keratinocyte abundance also contributes to UVB-induced tissue remodeling (**Fig. 6A**). We next examined whether these spatial changes were accompanied by transcriptional reprogramming of melanocytes. Heatmap analysis revealed that irradiated *Xpc*^−/−^ melanocytes displayed a transcriptional profile distinct from those of the other experimental groups across the basal, middle, and distal ear regions (**Fig. 6B**). These findings indicate that XPC deficiency markedly modifies the melanocyte transcriptional response to chronic UVB exposure across the longitudinal ear axis.

**Figure 6.**
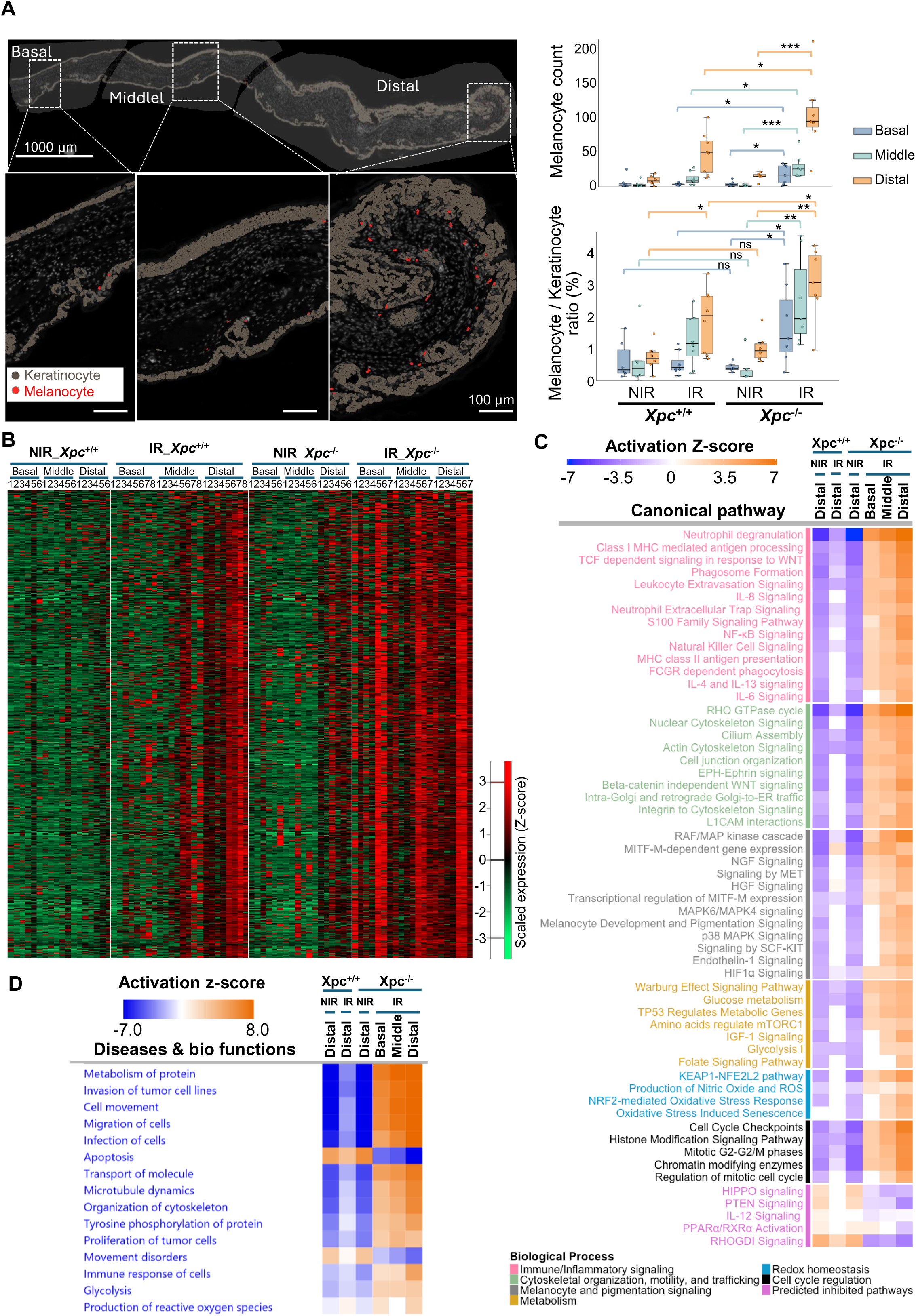
Spatial transcriptomics reveals melanocyte expansion and coordinated alterations in melanogenic, cytoskeletal, inflammatory, and metabolic programs in UVB-irradiated *Xpc*-deficient skin. All analyses were performed using Xenium spatial transcriptomic data from four experimental groups: *Xpc*^+/+^_NIR (n = 6), *Xpc*^+/+^_IR (n = 8), *Xpc*^−/−^_NIR (n = 6), and *Xpc*^−/−^_IR (n = 7), where n represents the number of mice. **(A)** Representative spatial maps showing melanocyte distribution along the longitudinal axis of a UVB-irradiated *Xpc*^−/−^ ear (left), with quantification of melanocyte abundance (top right) and the melanocyte-to-keratinocyte ratio (bottom right) across basal, middle, and distal ear regions under the four experimental conditions. Cell counts were determined from Xenium spatial transcriptomic data based on cell-type annotations. Statistical significance was assessed using Statistical significance was assessed using two-way ANOVA, with *P* < 0.05 considered statistically significant. *P < 0.05; **P < 0.01, *** P < 0.001. ns, not significant. **(B)** Heatmap showing melanocyte gene expression profiles across the four experimental conditions and three anatomical ear regions (basal, middle, and distal). Each column represents the aggregated transcript expression profile of melanocytes from one mouse within a defined anatomical region. The green-to-red color scale represents normalized, Z-score-transformed transcript abundance, with green indicating lower expression and red indicating higher expression. **(C)** Ingenuity Pathway Analysis (IPA) of canonical pathways associated with differentially expressed genes in melanocytes, highlighting alterations in immune and inflammatory signaling, cytoskeletal organization, melanocyte differentiation and pigmentation, metabolism, redox homeostasis, and cell cycle regulation. Orange and blue indicate predicted pathway activation and inhibition, respectively, based on IPA activation z-scores. Pathway labels are color-coded according to the biological processes indicated in the legend. **(D)** IPA Diseases and Bio Functions analysis of differentially expressed genes in melanocytes. Orange and blue indicate predicted activation and inhibition, respectively, based on IPA activation *z*-scores. NIR, nonirradiated; IR, UVB-irradiated.

Functional enrichment analysis using IPA further revealed coordinated changes in pathways associated with melanocyte development and pigmentation, together with MITF-dependent gene expression, cytoskeletal remodeling, glucose metabolism, inflammatory signaling, and redox homeostasis in irradiated *Xpc*^−/−^ melanocytes (**Fig. 6C and Fig. S6**). Consistent with these findings, IPA Diseases and Bio Functions analysis predicted increased cell movement and migration, immune responses, and reactive oxygen species production in irradiated *Xpc*^−/−^ melanocytes (**Fig. 6D**).

Together, these spatial and transcriptional analyses indicate that XPC deficiency enhances the UVB-induced expansion and redistribution of melanocytes and is associated with their coordinated transcriptional reprogramming toward a melanogenic, cytoskeletally remodeled, metabolically altered, and stress-responsive state.

### UVB induces a distinct pigmentation-supportive keratinocyte state in *Xpc*-deficient skin

Because keratinocytes are key regulators of melanocyte homeostasis and pigment transfer, we next investigated whether XPC deficiency alters keratinocyte states following chronic UVB exposure. Single-cell transcriptomic profiles of keratinocytes were extracted from the Xenium datasets and analyzed independently.

Heatmap analysis revealed a distinct transcriptional program in irradiated *Xpc*^−/−^ keratinocytes compared with the other experimental groups across the basal, middle, and distal ear regions (**Fig. 7A**). Functional enrichment analysis using IPA identified coordinated alterations in inflammatory signaling, pyrimidine salvage, glycolysis, tryptophan metabolism, pigmentation-supporting signaling, redox homeostasis, and cytoskeletal organization in irradiated *Xpc*^−/−^ keratinocytes (**Fig. 7B and Fig. S7A**). Conversely, PTEN, HIPPO, and adipogenesis-related pathways were predicted to be inhibited, further supporting extensive transcriptional and metabolic remodeling (**Fig. 7B**). PCA further identified genes contributing strongly to the separation of irradiated *Xpc*^−/−^ keratinocytes, including *Gjb2*, *Gjb6*, *Sprr2d/f*, *Cemip*, *Fscn1*, *Mcam*, *Tnc*, *Aldh1a3*, and *Ccl8*, collectively consistent with the emergence of a stress-responsive epidermal program involving inflammatory signaling, tissue remodeling, and cytoskeletal reorganization (**Fig. S7B**).

**Figure 7.**
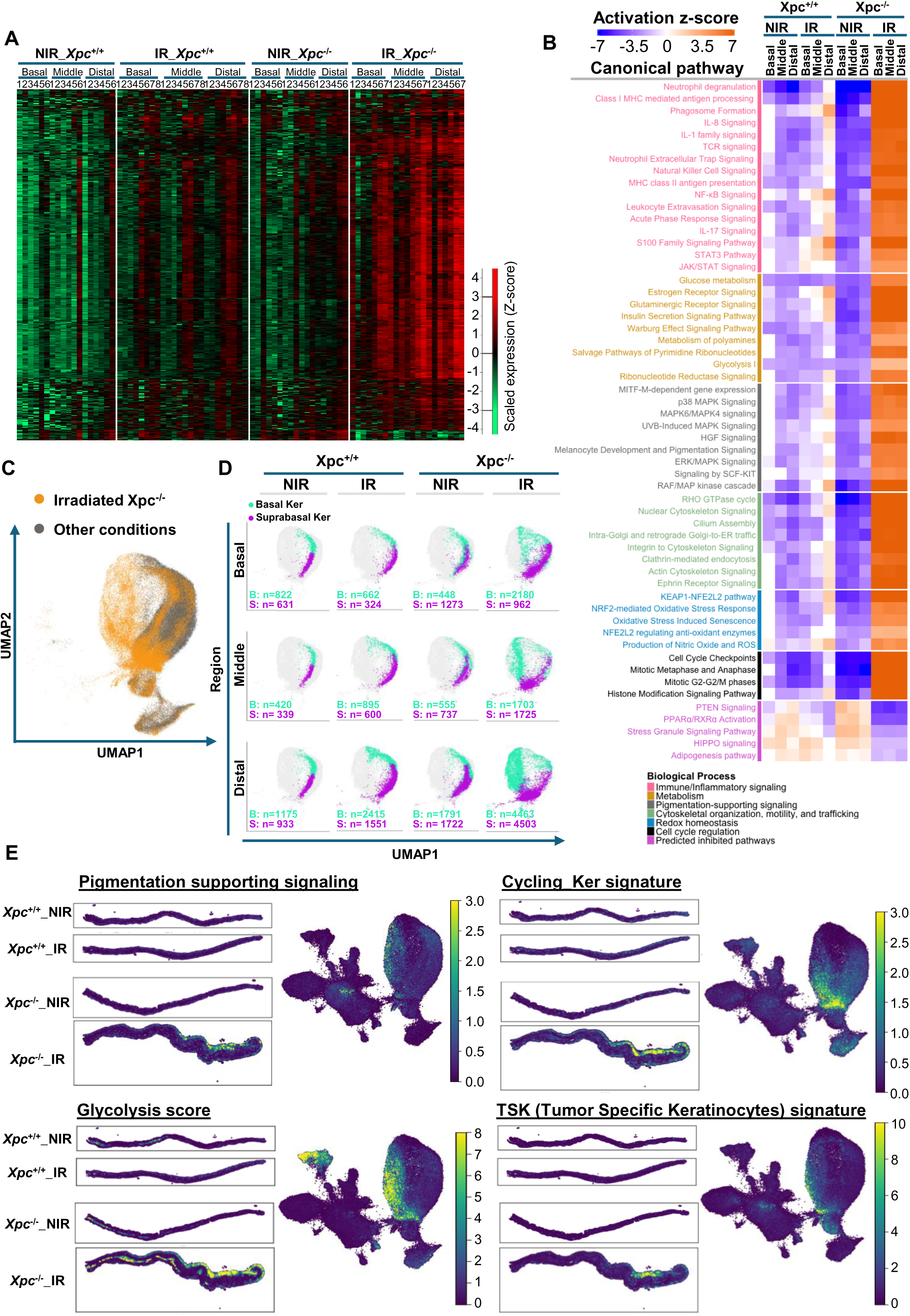
Spatial transcriptomics identifies a distinct stress-associated keratinocyte state with pigmentation-supporting features in UVB-irradiated Xpc-deficient skin. All analyses were performed using Xenium spatial transcriptomic data from four experimental groups: *Xpc*^+/+^_NIR (n = 6), *Xpc*^+/+^_IR (n = 8), *Xpc*^−/−^_NIR (n = 6), and *Xpc*^−/−^_IR (n = 7), where n represents the number of mice. (**A**) Heatmap showing keratinocyte gene expression profiles across the four experimental conditions and three anatomical ear regions (basal, middle, and distal). Each column represents the aggregated transcript expression profile of keratinocytes from one mouse within a defined anatomical region. The green-to-red color scale represents normalized, Z-score-transformed transcript abundance, with green indicating lower expression and red indicating higher expression. (**B**) Ingenuity Pathway Analysis (IPA) of canonical pathways associated with differentially expressed genes in keratinocytes. Orange and blue indicate predicted pathway activation and inhibition, respectively, based on IPA activation z-scores. Pathway labels are color-coded according to the biological processes indicated in the figure legend. (**C**) UMAP projection of the global keratinocyte transcriptomic landscape across the four experimental conditions. Keratinocytes from UVB-irradiated *Xpc*^−/−^ samples are highlighted in orange, whereas those from the other three experimental conditions are shown in gray, illustrating their distribution within the shared transcriptional space. (**D**) UMAP projections showing basal (B) and suprabasal (S) keratinocyte subpopulations across the three anatomical ear regions and four experimental conditions. Numbers indicate the corresponding cell counts for each keratinocyte subpopulation within each region and condition. (**E**) Spatial maps of gene-signature scores associated with pigmentation-supporting signaling, glycolysis, and transcriptional programs characteristic of cycling keratinocytes (Cycling_Ker), and tumor-specific keratinocytes (TSK) across the four experimental conditions. Scores were calculated from predefined gene sets and projected onto tissue coordinates, highlighting regional enrichment of these transcriptional programs, particularly in the middle and distal regions of UVB-irradiated *Xpc*^−/−^ skin. NIR, nonirradiated; IR, UVB-irradiated.

To further examine how the transcriptional changes associated with XPC deficiency were distributed across the global cellular landscape, we mapped cells from irradiated *Xpc*^−/−^ samples onto the UMAP containing all cells from the four experimental conditions (**Fig. 7C**). Cells from irradiated *Xpc*^−/−^ skin occupied distinct regions of the transcriptional landscape, with particularly prominent representation within the keratinocyte compartment. We therefore examined basal and suprabasal keratinocytes according to their anatomical location. Regional analysis revealed a progressive basal-to-distal transcriptional divergence that was particularly pronounced in irradiated *Xpc*^−/−^ skin (**Fig. 7D**). Whereas basal and suprabasal keratinocytes from the basal ear region of irradiated *Xpc*^−/−^ mice remained transcriptionally closer to their counterparts in the other experimental groups, those from the middle and, most prominently, distal ear regions progressively occupied distinct transcriptional states (**Fig. 7D**). Spatial signature scoring further revealed enrichment of pigmentation-supporting signalling, glycolytic metabolism, and proliferating/cycling keratinocyte programs within these regions (**Fig. 7E**). Notably, the tumor-specific keratinocyte (TSK) signature described by Ji et al. ^17^, a transcriptional state associated with epithelial plasticity and tissue remodeling, was also enriched, particularly toward the distal region (**Fig. 7E and Fig S5D**).

Together, these findings identify a spatially restricted keratinocyte state that emerges prominently in UVB-irradiated *Xpc*^−/−^ skin and is characterized by activation of DNA-damage and cell-cycle responses, cytoskeletal remodeling, inflammatory and stress signaling, and metabolic reprogramming. These changes were accompanied by increased cycling and glycolytic signatures and, notably, by enrichment of pigmentation-supporting and TSK-associated transcriptional programs, particularly in the distal ear, where melanocyte expansion and hyperpigmentation were most pronounced. This spatial and transcriptional association suggests that this keratinocyte state may contribute to a tissue microenvironment that supports persistent hyperpigmentation.

### Spatial transcriptomics identifies inflammatory and immunoregulatory immune niches associated with hyperpigmentation in UVB-irradiated *Xpc*-deficient skin

Because both melanocytes and keratinocytes exhibited prominent inflammatory transcriptional programs, we next investigated whether XPC deficiency alters immune-cell recruitment, composition, and spatial organization following chronic UVB exposure.

Immune-cell transcriptomic profiles were extracted from the Xenium datasets and analyzed independently. Differential expression analysis revealed a distinct transcriptional profile in immune cells from UVB-irradiated *Xpc*^−/−^ skin compared with the other experimental groups (**Fig. S8A**). IPA predicted broad immune activation involving neutrophil degranulation, phagosome formation and Fc receptor-mediated phagocytosis, cytokine and NF-κB signaling, antigen processing and presentation, and adaptive immune pathways including Th1 and T-cell signaling (**Fig. S8B**). IL-4/IL-13 signaling was also predicted to be activated, indicating a type-2 component within this broader inflammatory response, while interferon- and nucleic acid-sensing programs, including IFNα/β and cGAS–STING signaling, were also prominent (**Fig. S8B,C**). Consistent with these pathway-level changes, immune cells from irradiated *Xpc*^−/−^ skin showed increased expression of genes associated with myeloid activation (*Csf1r, Fcgr1, Fcgr4, Cd68,* and *Itgax*), interferon responses (*Ifi44, Ifit3, Isg15, Irf7,* and *Usp18*), leukocyte recruitment (*Ccl3, Ccl12, Cxcl10, Ccr1,* and *Ccr5*), and immune regulation (*Foxp3, Tigit, Pdcd1lg2,* and *Vsir*) (**Fig. S8C**). Alterations in redox homeostasis and glycolytic programs further indicated metabolic adaptation of the immune compartment (**Fig. S8C**). Accordingly, IPA Diseases and Bio Functions analysis predicted increased leukocyte development, migration, adhesion, extravasation, and phagocytic activity in irradiated *Xpc*^−/−^ skin (**Fig. S8D**).

To define the cellular composition underlying these transcriptional changes, immune cells were further resolved into distinct populations, including CCL8+ macrophages, tissue-resident macrophages, inflammatory macrophages, dendritic cells, mature migratory dendritic cells, monocytes, neutrophils, T cells/Tregs, and mast cells (**Fig. 8A and Fig. S9A**). Macrophage reclustering further resolved transcriptionally distinct tissue-resident, CCL8+, and inflammatory macrophage states (**Fig. S9**). CCL8+ macrophages were characterized by prominent chemokine, Fc receptor, and endolysosomal/phagocytic programs, whereas inflammatory macrophages displayed a distinct stress-responsive, glycolytic, cytoskeletal, and tissue-remodeling profile (**Fig. S9**).

**Figure 8.**
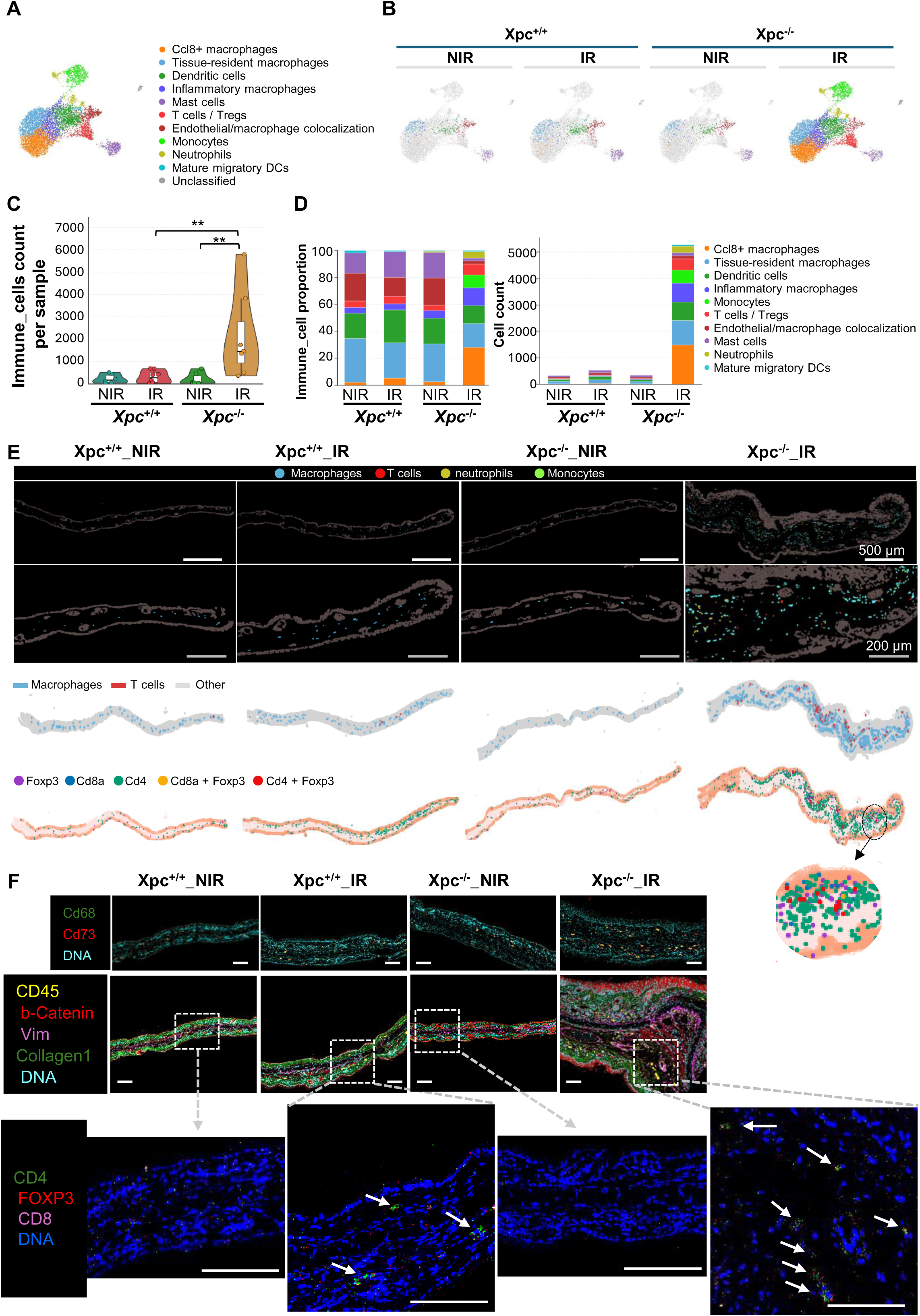
Spatial transcriptomics and Hyperion imaging mass cytometry reveal inflammatory immune niches associated with hyperpigmentation in UVB-irradiated *Xpc*-deficient skin. (**A-E**) Analyses of immune-cell populations using Xenium spatial transcriptomic data from four experimental groups: *Xpc*^+/+^_NIR (n = 6), *Xpc*^+/+^_IR (n = 8), *Xpc*^−/−^_NIR (n = 6), and *Xpc*^−/−^_IR (n = 7), where n represents the number of mice. (**A**) UMAP projection of immune-cell populations identified following extraction and reclustering of immune cells from the global Xenium dataset. Colors indicate the annotated immune-cell populations. (**B**) UMAP projections of immune cells split by experimental condition, illustrating their distribution across the four groups and the increased representation of immune cells from UVB-irradiated *Xpc*^−/−^ samples. (**C**) Violin plot showing immune-cell abundance across the four experimental conditions. Statistical significance was assessed using two-way ANOVA with * *P* < 0.05, and *** *P* < 0.001. (**D**) Bar plots showing the relative proportions (left) and absolute counts (right) of annotated immune-cell populations across the four experimental conditions. (**E**) Spatial distribution of immune-cell populations across the four experimental conditions. Upper panels show annotated immune cells projected onto Xenium tissue sections. Lower panels show the longitudinal distribution of macrophages and T cells (upper row), cells expressing *Cd4*, *Cd8a*, or *Foxp3* (lower row), along the ear axis. (**F**) Multiplex imaging mass cytometry (Hyperion) of immune-cell infiltration in ear skin across the four experimental conditions. Top panels show representative images of CD68 (green), CD73 (red), and DNA (cyan), highlighting increased accumulation of CD68-positive myeloid cells in UVB-irradiated *Xpc*^−/−^ skin. Middle panels show CD45 (yellow), β-catenin (red), collagen I (green), vimentin (magenta), and DNA (cyan). Dashed boxes indicate regions of CD45-positive immune-cell infiltration shown at higher magnification in the bottom panels. Bottom panels show CD4 (green), FOXP3 (red), CD8 (magenta), and DNA (cyan), illustrating the presence of CD4-positive and less frequent CD8-positive cells within the corresponding immune-cell infiltrates. Scale bars, 100 µm. NIR, nonirradiated; IR, UVB-irradiated.

Comparison across experimental conditions revealed a pronounced enrichment of immune cells in UVB-irradiated *Xpc*^−/−^ skin (**Fig. 8B,C**). This expansion was not restricted to a single immune population. Whereas the relatively sparse immune compartment in the other conditions was predominantly composed of macrophages, irradiated *Xpc*^−/−^ skin displayed expansion of multiple macrophage states together with increased monocytes, neutrophils, dendritic cells, and T cells (**Fig. 8D**).

Spatial mapping revealed that this immune expansion was regionally organized, with macrophages and T cells preferentially accumulating in the middle and distal regions of irradiated *Xpc*^−/−^ ears (**Fig. 8E**), coinciding with the regions displaying the greatest melanocyte expansion and pigmentation. T-cell annotation indicated a predominance of CD4+ cells, including a population co-expressing *Cd4* and *Foxp3* (**Fig. 8E**). Multiplex imaging mass cytometry (Hyperion) performed on the same tissue specimens confirmed the increased immune infiltration at the protein level. Irradiated *Xpc*^−/−^ skin showed marked accumulation of CD68+ myeloid cells and CD45+ leukocytes compared with the other experimental groups, whereas immune infiltration in irradiated *Xpc*+/+ skin was more limited (**Fig. 8F**). Within CD45+ infiltrated regions, lymphocytic infiltration was predominantly composed of CD4+ cells, with comparatively few CD8+ cells, and numerous CD4+ cells co-expressed FOXP3 (**Fig. 8F**).

Together, these findings reveal a marked expansion and transcriptional activation of the immune compartment in UVB-irradiated *Xpc*^−/−^ skin, dominated by macrophage/myeloid populations and characterized by a broad inflammatory response in which type-2 features coexist with interferon, Th1, phagocytic, and immunoregulatory programs. These immune populations form spatially organized inflammatory and immunoregulatory niches concentrated in regions of pronounced melanocyte expansion and pigmentation. Their spatial convergence with the keratinocyte and melanocyte states identified above supports coordinated multicellular remodeling of the cutaneous microenvironment during persistent UVB-induced hyperpigmentation.

### Human XP-C skin and XPC-deficient melanocytes exhibit pigmentation-associated alterations overlapping with those observed in UVB-irradiated XPC-deficient mice

To determine whether the molecular and cellular alterations identified in the XPC-deficient mouse model are recapitulated in human disease, we next analyzed paired hypo- and hyperpigmented skin regions obtained from three XP-C patients (**Supplementary Table 1**). Histological examination revealed striking pigmentary heterogeneity within individual biopsies, characterized by sharply demarcated hyperpigmented and hypopigmented areas (**Fig. 9A**). Compared with healthy skin, hyperpigmented lesions displayed markedly increased melanin deposition throughout the basal epidermis, whereas hypopigmented regions exhibited a profound reduction in pigment content (**Fig. 9A**). Interestingly, focal clusters of keratinocytes retaining supranuclear melanin caps were frequently observed within hyperpigmented regions, consistent with enhanced melanosome transfer and retention. Immunofluorescence analyses further demonstrated that melanocyte density was significantly increased within hyperpigmented regions (**Fig. 9B**). In contrast, melanocytes were scarce within hypopigmented areas, although residual TRP-1-positive cells remained detectable in some lesions. In addition, melanocytes occasionally appeared partially detached from the basement membrane in hypopigmented areas, suggesting altered melanocyte adhesion or tissue organization (**Fig. 9B**). Because our mouse studies identified alterations in melanosome trafficking, we next examined the expression of the melanosome transport regulator RAB27A together with the melanocyte marker TRP-1. In healthy skin and hyperpigmented XP-C lesions, RAB27A strongly co-localized with TRP-1-positive melanocytes, whereas RAB27A-positive melanocytes were virtually absent from hypopigmented regions (**Fig. 9C**). Thus, hyperpigmented XP-C lesions are characterized not only by increased melanocyte abundance and melanin deposition but also by activation of the melanosome transport machinery. Together, these findings demonstrate that human XP-C pigmentary lesions closely phenocopy the UVB-induced phenotype observed in XPC-deficient mice, supporting the translational relevance of the mouse model and suggesting that dysregulated melanocyte distribution, melanosome trafficking and pigment transfer represent conserved features of XPC deficiency.

**Figure 9.**
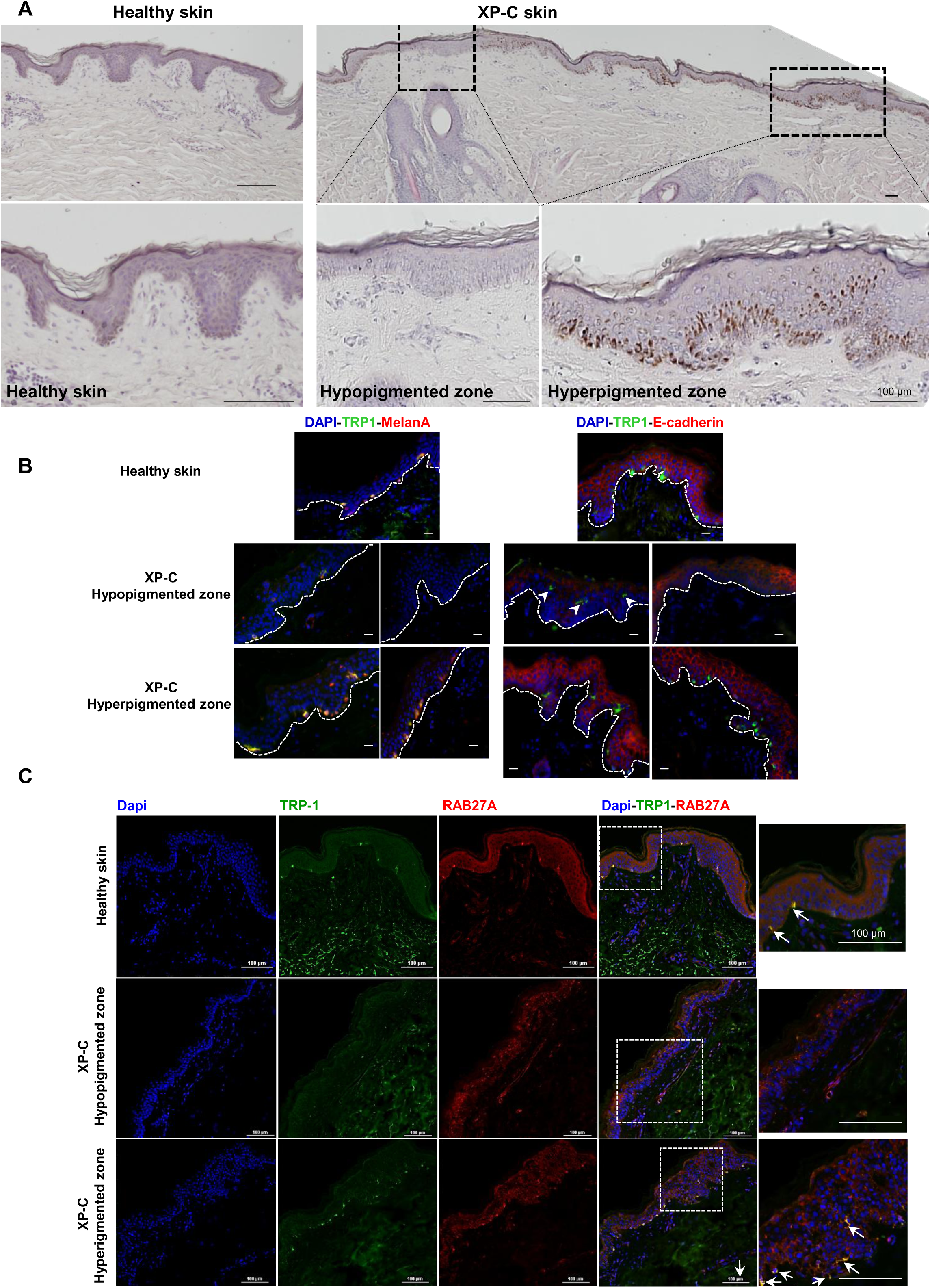
Human XP-C skin exhibits alterations in melanocyte distribution and melanosome-associated transport machinery across hyperpigmented and hypopigmented regions. (**A**) Representative brightfield images of healthy control skin (left) and XP-C skin (right), illustrating the coexistence of hypo- and hyperpigmented regions within the same biopsy. Higher-magnification views of the dashed boxes highlight reduced melanin deposition in hypopigmented areas and pronounced pigment accumulation in the basal epidermis of adjacent hyperpigmented regions. Scale bars, 100 μm. **(B)** Immunofluorescence characterization of healthy control skin and hypopigmented and hyperpigmented regions from an XP-C patient. Left panels show melanocyte distribution using the melanocytic markers TRP-1 (green) and Melan-A (red). Right panels show TRP-1 (green) and E-cadherin (red), highlighting epidermal architecture and melanocyte localization. Hyperpigmented regions exhibit increased melanocyte abundance, whereas hypopigmented regions contain relatively few detectable melanocytes. White dashed lines delineate the dermal–epidermal junction, and arrowheads indicate representative melanocytes with altered positioning relative to this boundary. Nuclei are counterstained with DAPI (blue). Scale bars, 100 μm. **(C)** Immunofluorescence analysis of RAB27A (red), a regulator of melanosome transport, and the melanocyte marker TRP-1 (green) in healthy control skin and hypopigmented and hyperpigmented regions from an XP-C patient. Merged images show overlapping RAB27A and TRP-1 signals in healthy skin and hyperpigmented XP-C regions, whereas cells coexpressing these markers are rarely detected in hypopigmented regions. Dashed boxes indicate areas shown at higher magnification in the right panels. White arrows highlight representative regions of RAB27A and TRP-1 signal overlap within melanocytes. Nuclei are counterstained with DAPI (blue). Scale bars, 100 μm..

To determine whether the molecular alterations identified in XPC-deficient mice were also present in human disease, paired hypo- and hyperpigmented regions from three XP-C patients were subjected to label-free quantitative proteomic analysis. PCA analysis revealed a clear segregation between hypo- and hyperpigmented samples, indicating distinct proteomic profiles associated with pigmentation status (**Fig. 10A**). Consistently, unsupervised hierarchical clustering separated samples according to their pigmentation status, revealing relative similarities in proteomic profiles within each group despite their different patient origins (**Fig. 10B**). Functional enrichment analysis using IPA identified alterations in inflammatory and immune signaling pathways, cytoskeletal remodeling, MAP kinase signaling, JAK–STAT signaling, and pathways associated with oxidative stress responses and cellular redox homeostasis in hyperpigmented regions (**Fig. 10C**). Notably, these biological processes were also affected in UVB-irradiated *Xpc*^−/−^ mouse skin, particularly in regions exhibiting pronounced hyperpigmentation.

**Figure 10.**
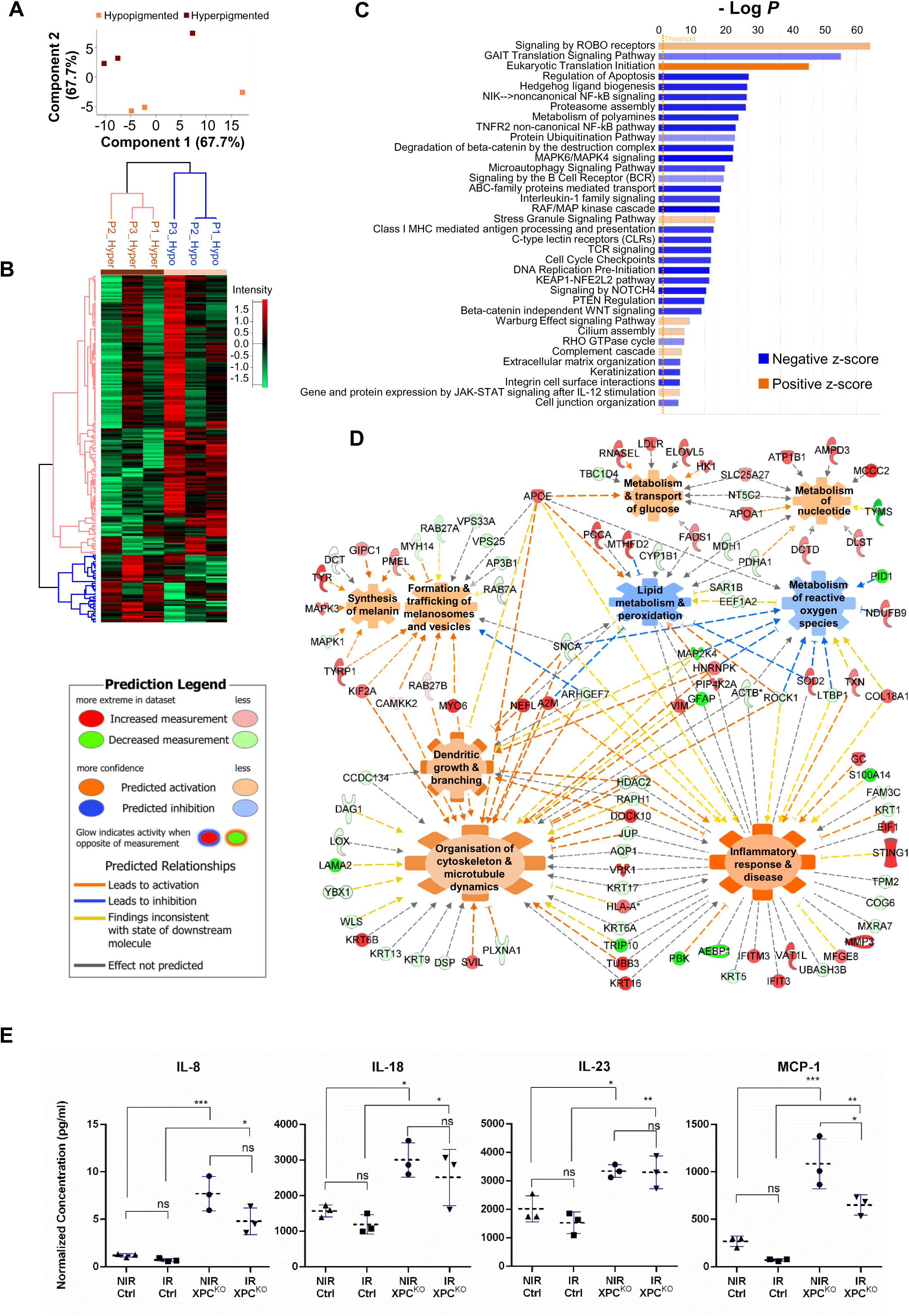
Proteomic profiling and functional analyses reveal inflammatory, cytoskeletal, and metabolic alterations in human XP-C skin and XPC-deficient melanocytes. (**A-C**) hypo- and hyperpigmented skin regions of three XP-C patients were subjected to label-free quantitative proteomic. **(A)** Principal component analysis (PCA) demonstrates clear segregation of hyperpigmented (brown) and hypopigmented (orange) samples based on their global protein expression profiles. **(B)** Unsupervised hierarchical clustering of differentially expressed proteins (rows) across paired hypo- and hyperpigmented XP-C skin samples (columns). The green-to-red color scale represents relative protein abundance following normalization and Z-score transformation, with green indicating lower abundance and red indicating higher abundance. Hierarchical clustering was performed using Spearman correlation and complete linkage in Perseus. **(C)** Canonical pathway enrichment analysis generated using Ingenuity Pathway Analysis (IPA). Pathways are ranked according to statistical significance (−log10 *P*). Orange bars indicate predicted pathway activation (positive activation z-score), whereas blue bars indicate predicted inhibition (negative activation z-score). The dashed vertical line represents the significance threshold. **(D)** CRISPR/Cas9-generated XPC knockout (XPC^KO^) and control primary human melanocytes were subjected to label-free quantitative proteomic analysis (n = 5 independent biological replicates per group). The output of differentially expressed proteins on the indicated biological functions were analyzed using IPA. Protein nodes are colored according to measured abundance (red, increased; green, decreased), whereas biological functions are colored according to their predicted activation state (orange, activated; blue, inhibited). **(E)** Multiplex immunoassay quantification of cytokines secreted by CRISPR/Cas9-generated control and XPC knockout primary human melanocytes cultured under non-irradiated (NIR) or UVB-irradiated (IR) conditions. Secreted concentrations of IL-8, IL-18, IL-23, and MCP-1 are shown. Data are the mean ± SD of three independent biological replicates per condition. Statistical significance was determined using one-way ANOVA followed by Tukey’s post hoc test. \**P* < 0.05, **P < 0.01, \*\*\**P* < 0.001.

Further mechanistic network analyses highlighted interconnected molecular networks associated with inflammatory and immune responses (**Fig. S10A**), epidermal differentiation and keratinization (**Fig. S10B**), lipid metabolism and MAP kinase signaling (**Fig. S10C**), and cytoskeletal organization, cell adhesion, and mechanotransduction (**Fig. S10D**). In particular, TNF-centered inflammatory networks involved components of NF-κB signaling, SRC-family kinases, and immune receptor signaling (**Fig. S10A**), whereas additional networks highlighted alterations in keratinocyte differentiation, lipid metabolism, MAP kinase signaling, and actomyosin-associated cytoskeletal organization (**Fig. S10B–D**). Together, these analyses suggest that hyperpigmented XP-C skin is characterized by coordinated alterations in inflammatory signaling, epidermal organization, and metabolic and cytoskeletal processes, consistent with several molecular features identified in UVB-irradiated *Xpc*^−/−^ mouse skin.

To further investigate the contribution of XPC deficiency to melanocyte-associated molecular alterations, we generated stable XPC-deficient primary human melanocytes using CRISPR/Cas9 genome editing, as previously described ^18^. Proteomic analysis followed by IPA identified molecular networks associated with melanin synthesis, melanosome trafficking, dendritic growth, cytoskeletal organization, inflammatory responses, and metabolic regulation (**Fig. 10D**). These findings suggest that XPC deficiency is associated with alterations in multiple melanocyte-related biological programs, independently of the surrounding skin microenvironment.

Consistent with changes in inflammatory signaling, multiplex cytokine analysis revealed increased secretion of several inflammatory mediators, including IL-8, IL-18, IL-23, and MCP-1, in nonirradiated XPC-deficient melanocytes compared with controls (**Fig. 10E**). UVB irradiation did not consistently further increase cytokine secretion in XPC-deficient cells. Together, these findings demonstrate that XPC deficiency is associated with a cell-intrinsic pro-inflammatory secretory phenotype that is evident without UVB irradiation and remains detectable following UVB exposure.

Together, these findings reveal alterations in melanocyte distribution, pigmentation-associated processes, and inflammatory, cytoskeletal, and metabolic programs in human XP-C skin and XPC-deficient melanocytes. The overlap between these alterations and those identified in UVB-irradiated *Xpc*^−/−^ mouse skin supports the translational relevance of our experimental model.

## Discussion

Our study reframes persistent DNA damage not as a passive lesion awaiting repair, but as an active signal capable of orchestrating tissue-wide adaptation. This concept builds on emerging evidence that unresolved genomic injury can induce senescence-associated secretory programs and remodel neighboring immune and stromal compartments ^19^, while DNA damage signaling can rewire cellular metabolism far beyond the nucleus ^20–22^. Whereas previous studies have largely focused on cell-autonomous or pathway-specific responses to genotoxic stress, our findings show that persistent DNA damage can coordinate the behavior of multiple cell populations within an intact tissue. By integrating spatial transcriptomics with proteomics, metabolomics, kinome profiling and functional analyses, we demonstrate that XPC deficiency drives coordinated reprogramming of melanocytes, keratinocytes and immune cells toward a stress-responsive immunometabolic tissue state whose cumulative output is persistent hyperpigmentation. These findings extend previous reports that XPC-deficient cells can establish a permissive local microenvironment ^18,23,24^ by showing that this response is not restricted to a single cell lineage, but instead involves multicellular remodeling across the cutaneous microenvironment. Chronic UV-induced pigmentation therefore emerges as a systems-level adaptation to persistent genotoxic stress, a paradigm that may extend to other DNA repair-deficient and aging-associated skin disorders.

Current models of UV-induced pigmentation largely emphasize paracrine communication between keratinocytes and melanocytes through pathways such as p53–POMC–α-MSH or KITLG–KIT signaling ^25,26^. While these mechanisms undoubtedly contribute to physiological tanning, they do not fully explain the persistent architectural and cellular remodeling observed following chronic UV exposure. Using XPC deficiency as a model of defective genome surveillance, our study identifies persistent DNA damage itself as a plausible upstream driver of these long-term adaptations. The remarkable concordance between the murine phenotype and XP-C patient biopsies further supports the concept that unrepaired DNA lesions initiate sustained tissue remodeling extending well beyond the DNA repair machinery.

One of the most consistent features emerging from our multi-omics analyses was extensive remodeling of cellular redox homeostasis. Increased glutathione disulfide, depletion of taurine and hypotaurine, altered nicotinamide metabolism, and activation of glutathione-associated pathways collectively indicate sustained oxidative stress in chronically UVB-irradiated Xpc-deficient skin. These findings are consistent with previous studies showing that UVB exposure disrupts glutathione redox balance and cellular redox homeostasis in the skin ^27^. These observed metabolic alterations were accompanied by induction of antioxidant enzymes and redox-responsive signaling pathways, suggesting that oxidative stress is not simply a downstream consequence of defective DNA repair but an integral component of the adaptive response to persistent photodamage. This interpretation is supported by previous studies implicating oxidative stress in XPC pathophysiology ^10,21,22^, melanocyte activation ^28^, and the pathogenesis of solar lentigo, in which oxidative stress-associated molecular signatures have been identified in human lesions ^29^. Our findings extend these observations by placing redox dysregulation within a broader multicellular framework, in which coordinated oxidative-stress responses across melanocytes, keratinocytes and immune cells contribute to sustain pigmentary remodeling.

One of the major conceptual implications of this study is that persistent UV-induced DNA damage does not selectively alter melanocytes but is associated with coordinated reprogramming of multiple cutaneous cell populations. Spatial transcriptomics revealed concurrent changes in melanocytes, keratinocytes, and infiltrating immune cells, involving inflammatory, metabolic, and cytoskeletal programs that spatially converged in regions of pronounced pigmentation. These findings support a model in which the persistent pigmentary phenotype of XPC deficiency emerges within a multicellular tissue state rather than from melanocyte-intrinsic alterations alone. Clinical observations in patients with XP provide intriguing support for a role of the local immune environment in modulating this pigmentary state. Topical treatment with the immune-response modifier imiquimod has been reported to improve background hyperpigmentation and chronic actinic changes in XP, including long-lasting improvement of freckling and background hyperpigmentation in treated fields^30–32^. Although the mechanisms underlying these pigmentary effects remain uncertain, they demonstrate that at least some XP-associated hyperpigmentation is reversible despite persistence of the underlying DNA-repair defect. This observation is consistent with our finding that enhanced melanocyte activity occurs within a profoundly remodeled keratinocyte and immune microenvironment, and raises the possibility that altering local tissue signaling can modify the pigmentary phenotype.

Further support for this concept comes from a recent report of an XPC-deficient patient with advanced cSCC treated with the PD-1 inhibitor pembrolizumab. In addition to sustained tumor regression, treatment was accompanied by rapid and persistent fading of the patient’s widespread lentigines, whereas pre-existing vitiligo, melanocytic nevi, and seborrheic keratoses remained unchanged ^33^. Interestingly, lentigine regression has not been observed uniformly in all pembrolizumab-treated patients with XP and there are the cases in which tumor response occurred without fading of lentigines ^34^. It has been proposed that the pre-existing immune context, including vitiligo in their patient, may have amplified immune recognition of UV-damaged pigmented cells ^33^. Together, these clinical observations and our spatial data support the concept that XP-associated pigmentation is not simply a fixed consequence of melanocyte dysfunction, but a dynamic phenotype influenced by interactions among persistent UV damage, keratinocyte states, melanocyte activity, and the local immune microenvironment.

Our findings further identify cytoskeletal remodeling as a major determinant of chronic hyperpigmentation. Proteomic analyses, kinase profiling and spatial transcriptomics consistently revealed alterations in pathways controlling actin dynamics, integrin signaling and cytoskeletal organization, in line with previous studies demonstrating that cytoskeletal machinery governs melanocyte morphology, dendrite formation and pigment distribution ^35–37^. These molecular changes were accompanied by increased melanocyte surface area and dendricity, altered melanosome biogenesis, and markedly enhanced melanosome transfer to keratinocytes, processes that rely on coordinated actin- and microtubule-dependent trafficking ^35^. Together, these findings suggest that cytoskeletal reorganization provides a mechanistic link between persistent DNA damage and sustained pigment redistribution within the epidermis, by coupling melanocyte morphological plasticity to enhanced intracellular melanosome trafficking and transfer to neighboring keratinocytes.

A second major theme emerging from our study is the extensive metabolic adaptation associated with chronic UV exposure. Independent proteomic, metabolomic and kinome analyses converged on enhanced glycolysis, nucleotide biosynthesis and redox metabolism, together with activation of pathways supporting increased biosynthetic demand and oxidative-stress adaptation. This profile is consistent with the compartmentalized and energy-intensive nature of melanin synthesis and melanosome maturation described in melanocyte metabolism ^38^. Such metabolic remodeling likely supports not only sustained melanogenesis, but also the energetic and biosynthetic demands imposed by cytokine production, cellular proliferation and tissue remodeling. Importantly, metabolic reprogramming was not restricted to melanocytes but was also evident across keratinocyte and immune-cell compartments. This convergence echoes recent evidence that glycolytic reprogramming can function as a shared metabolic hub coordinating epithelial and immune responses during chronic skin inflammation ^39^. Together, these findings support a tissue-wide immunometabolic adaptation to persistent photodamage rather than a melanocyte-autonomous metabolic response.

Although XP-C is a rare inherited disorder, our findings may have broader implications for common acquired pigmentary disorders. Several hallmarks identified here, including altered melanocyte redistribution, increased eumelanin production, enhanced melanosome transfer, inflammatory remodeling and metabolic adaptation, have individually been reported in solar lentigo ^29^, chronic photoaging and post-inflammatory hyperpigmentation ^40^. Our study integrates these observations into a unified mechanistic framework and suggests that persistent DNA damage may represent a common upstream driver of chronic pigmentary remodeling. In this context, XP-C may be viewed as an accelerated model that reveals mechanisms otherwise developing over decades of cumulative sun exposure.

One notable difference between our mouse model and the human XP-C phenotype is the absence of obvious hypopigmented lesions following chronic UVB exposure. Several non-mutually exclusive explanations may account for this discrepancy. First, the duration of UVB exposure in our model was intentionally limited to capture early adaptive responses before the onset of severe tissue degeneration or carcinogenesis, whereas human XP-C patients accumulate decades of repeated solar exposure. Second, murine and human skin differ substantially in melanocyte distribution, with adult mice containing numerous dermal melanocytes that are largely absent from normal human interfollicular epidermis. The persistence and recruitment of these dermal melanocytes may partially compensate for local melanocyte depletion and favor a predominantly hyperpigmented phenotype. Finally, hypopigmented macules in XP patients may result from progressive melanocyte exhaustion, stem cell depletion or irreversible loss of pigment-producing cells following repeated cycles of DNA damage and inflammation, processes that are unlikely to be fully reproduced during the relatively short experimental period used in this study. Long-term irradiation models will therefore be required to determine whether the pro-pigmentary tissue state identified here eventually progresses toward the mixed hypo- and hyperpigmented phenotype characteristic of human disease.

### Strengths, limitations and conclusion

The principal strength of this study lies in the integration of complementary approaches spanning ultrastructural imaging, functional assays, quantitative multi-omics and spatial transcriptomics, together with complementary validation in both mouse and human systems. Although future studies will be required to dissect the precise causal interactions among keratinocytes, melanocytes and immune cells, our findings consistently converge on a common biological model. Rather than representing a purely melanocyte-autonomous response, chronic UV-induced pigmentation emerges as a multicellular adaptive process orchestrated by persistent DNA damage through coordinated oxidative, inflammatory, metabolic and cytoskeletal remodeling of the cutaneous microenvironment. These findings redefine XPC as a regulator of tissue adaptation to chronic UV stress and provide a mechanistic framework that may extend beyond XP-C to common UV-associated pigmentary disorders.

## Acknowledgments

We acknowledge the Biochemistry and Biophysics Platform (BioProt) of the Bordeaux Neurocampus for access to the PamStation (PamGene analysis), supported by the GPR BRAIN 2030 with the help of M. David and J. Ezan. Imaging was performed on the Bordeaux Imaging Center, member of the FranceBioImaging national infrastructure (ANR-10-INBS-04).

We acknowledge the SIMCAT facility of Montpellier Cancer Research Institute for technical and scientific expertise. The authors thank the staff of the SAM Metabolic Analyses Platform at TBMCore (University of Bordeaux, CNRS UAR 3427, INSERM US05) for their valuable assistance and expertise in the acquisition and analysis of metabolomics data. We also gratefully acknowledge Mr. Sebastien Tarrit (UMR 1240 Inserm/UCA) for his valuable technical assistance with melanin quantification.

## Ethics approval and informed consent

Human skin samples were obtained from three patients with XP-C. Written informed consent for participation and publication was obtained from the patients or their legal guardians, as applicable. The study was approved by the local ethics committee of the University Hospital of Bordeaux, France (CE-GR-2017/006).

---

**Supplementary Table 1.** Clinical and pathological characteristics of XP-C patients included in this study.

| **Patient ID** | **Sex** | **XPC mutation** | **Age at diagnosis (months)** | **Age at biopsy (years)** | **Biopsy site** |
|---|---|---|---|---|---|
| **P1** | Male | XPC c.1243C>T/p.Arg415X;c.1704T>A/p.Tyr568X | 18 | 23 | Forearm |
| **P2** | Male | XPC c.1243C>T/p.Arg415X;c.1704T>A/p.Tyr568X | 18 | 23 | Forearm |
| **P3** | Male | XPC (c.1639delTG/p.Val548Alafs*25) hmz | 24 | 15 | Forearm |

Paired hypo- and hyperpigmented skin biopsies were obtained from each XP-C patient and processed for histological, immunofluorescence and proteomic analyses, as indicated.

**Supplementary Table 2.** Antibody panel used for Hyperion imaging mass cytometry.

| Target | Clone | Provider | Tag | Dilution1 |
|---|---|---|---|---|
| Iba1 | EPR16588 | Abcam | **113Cd** | 1000 |
| Tyrosinase | T311 | SantaCruz | **141Pr** | 150 |
| SOX10 | EPR4007 | Abcam | **142Nd** | 100 |
| Vimentin | D21H3 | StandardBiotools | **143Nd** | 1000 |
| B220 | RA3-6B2 | StandardBiotools | **144Nd** | 800 |
| NCR1 | EPR23097-35 | Abcam | **145Nd** | 100 |
| E-Cadherin | Polyclonal | ProteinTech | **146Nd** | 100 |
| GranzymeB | D2H2F | CellSignalingTechnologies | **147Sm** | 200 |
| Pan CK | C11 | StandardBiotools | **148Nd** | 100 |
| Pan CK | AE1/AE3 | SantaCruz | **148Nd** | 100 |
| MLANA | Rmlana/788 | NovusBio | **149Sm** | 200 |
| PD-L1 | 2B11D11 | ProteinTech | **150Nd** | 100 |
| CD31 | EPR17259 | Abcam | **151Eu** | 200 |
| CD3 | CD3-12 | Abcam | **152Sm** | 200 |
| CD8a | EPR21769 | Abcam | **153Eu** | 100 |
| CD11c | D1V9Y | CellSignalingTechnologies | **154Sm** | 200 |
| RORgt | EPR20006 | Abcam | **155Gd** | 200 |
| CD80 | Polyclonal | Abcam | **156Gd** | 400 |
| Foxp3 | D6O8R | CellSignalingTechnologies | **158Gd** | 400 |
| CD68 | Polyclonal | ProteinTech | **159Tb** | 1000 |
| CD208/Lamp3 | Polyclonal | ProteinTech | **160Gd** | 400 |
| PD1 | D7D5W | CellSignalingTechnologies | **161Dy** | 100 |
| PNAD-biotin | MECA-79 | Biolegend | **162Dy** | 1000 |
| CD4 | BLR167J | Bethyl | **163Dy** | 200 |
| CD45 | Polyclonal | ProteinTech | **164Dy** | 100 |
| B-Catenin | D13A1 | StandardBiotools | **165Ho** | 400 |
| MHC-II | M5/114.15.2 | Novus | **166Er** | 1000 |
| GATA3 | EPR16651 | Abcam | **167Er** | 100 |
| Ki67 | B56 | StandardBiotools | **168Er** | 500 |
| Collagen type I | Polyclonal | StandardBiotools | **169Tm** | 200 |
| F4/80 | D2S9R | CellSignalingTechnologies | **170Er** | 200 |
| Lyve1 | Polyclonal | Abcam | **171Er** | 100 |
| Caspase-3 | 5A1E | StandardBiotools | **172Yb** | 200 |
| CD103 | EPR22590-27 | Abcam | **174Yb** | 100 |
| CD73 | Polyclonal | ProteinTech | **175Lu** | 200 |
| CD39 | EPR20627 | Abcam | **176Yb** | 100 |
| MPO | EPR20257 | Abcam | **194Pt** | 1000 |
| Alpha-SMA | 1A4 | StandardBiotools | **209Bi** | 1000 |

## Supplementary figure legends

**Supplementary Figure S1.**
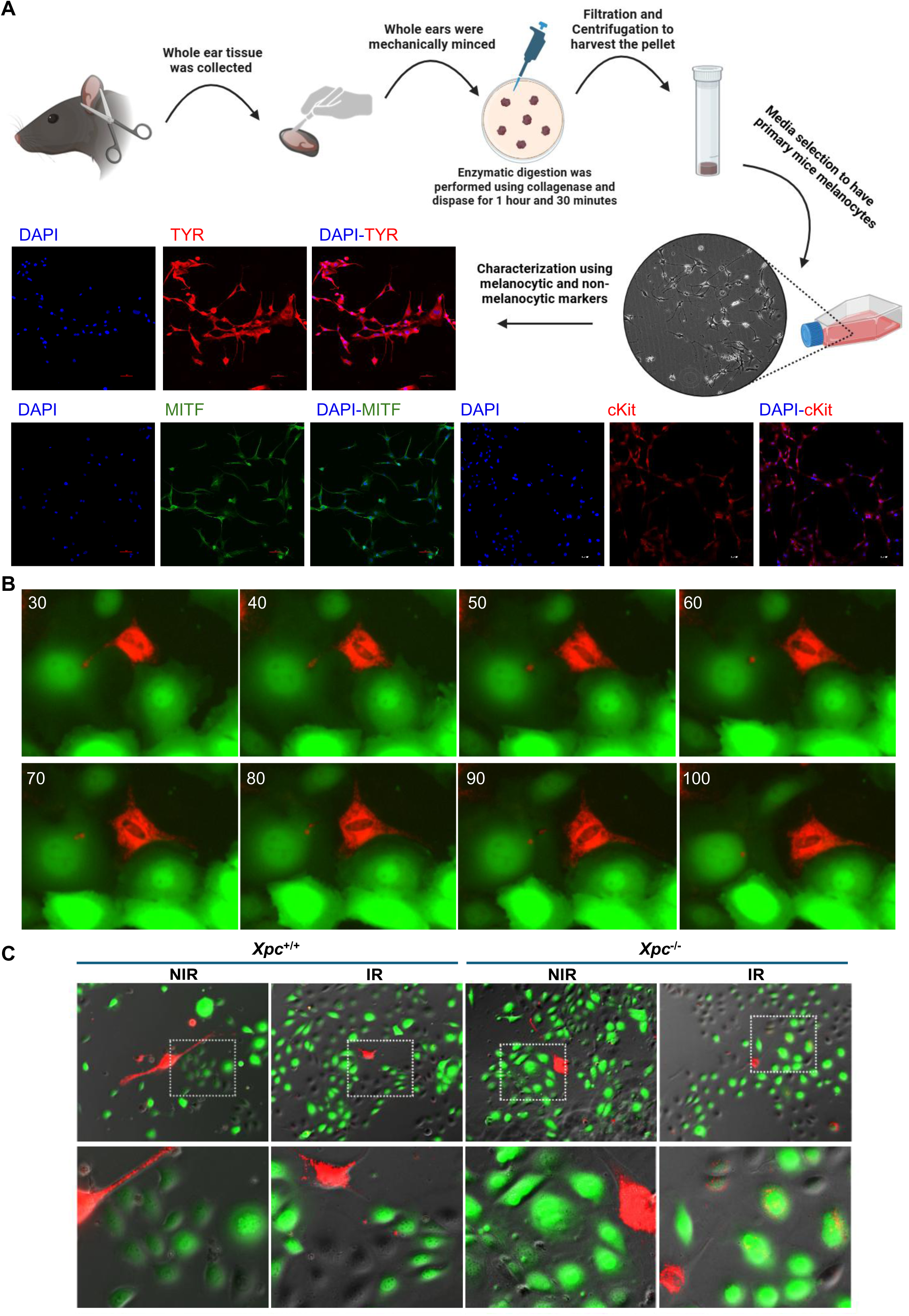
XPC deficiency enhances UVB-induced melanosome transfer from melanocytes to keratinocytes. (**A**) Experimental workflow for the isolation and characterization of primary mouse melanocytes. Ear skin was mechanically dissociated and enzymatically digested using collagenase and dispase, followed by filtration, centrifugation, and culture under melanocyte-selective conditions. Immunofluorescence staining confirmed melanocyte identity using the melanocytic markers TYR, MITF, and c-KIT. Nuclei were counterstained with DAPI. **(B–C)** Visualization of melanosome transfer in melanocyte–keratinocyte co-cultures. Primary *Xpc*^+/+^ or *Xpc*^−/−^ melanocytes were transduced with MelanoTag (mScarlet-GPR143) to fluorescently label melanosomes and co-cultured with GFP-expressing mouse keratinocytes at a 1:10 ratio. Co-cultures were either sham-irradiated (NIR) or exposed to two UVB irradiations (5 mJ/cm^2^ each) separated by 48 h. Time-lapse fluorescence imaging was initiated immediately after the second UVB exposure to monitor melanosome transfer dynamics. **(B)** Representative time-lapse fluorescence images showing the transfer of MelanoTag-positive melanosomes from a UVB-irradiated *Xpc*^−/−^ melanocyte into an adjacent GFP-expressing keratinocyte. Images were acquired at 10-min intervals over a 70-min period. Time labels indicate minutes elapsed from the beginning of imaging. **(C)** Representative phase-contrast and fluorescence images of co-cultures acquired 5 h after the second irradiation. Lower panels show higher-magnification views of the boxed regions, highlighting transferred MelanoTag-positive melanosomes within GFP-positive keratinocytes. NIR, nonirradiated; IR, UVB-irradiated.

**Supplementary Figure S2.**
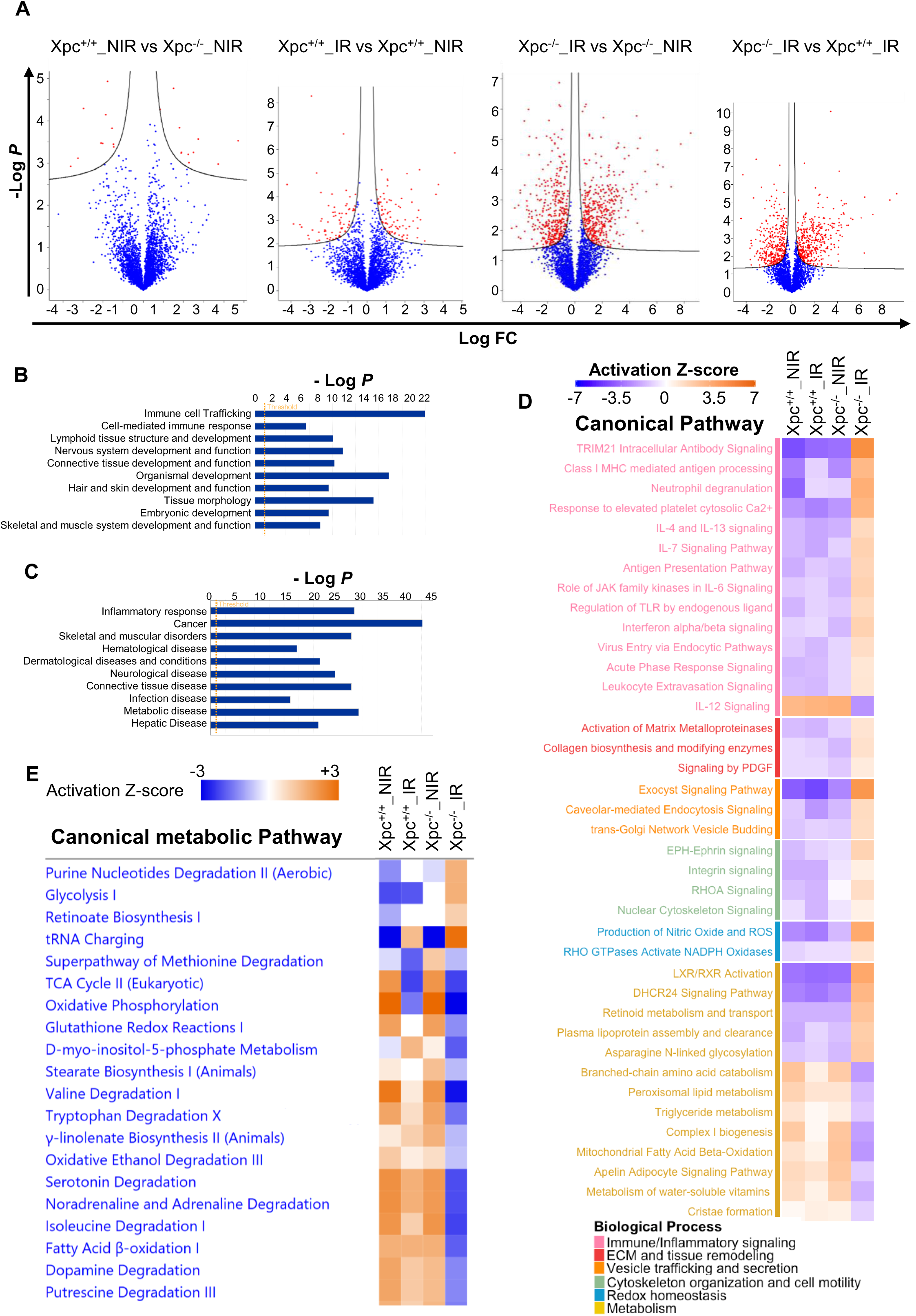
Proteomic profiling reveals inflammatory activation, metabolism rewiring, oxidative stress responses and cytoskeletal remodeling in UVB-irradiated Xpc-deficient skin. (**A**) Volcano plots showing global protein expression changes across the indicated pairwise comparisons. The x-axis represents log2 fold change and the y-axis represents −log10 P value. Significantly altered proteins are highlighted in red. Differentially expressed proteins were identified using permutation-based false discovery rate (FDR) correction (FDR = 0.05). (**B**) Functional enrichment analysis of differentially expressed proteins using Ingenuity Pathway Analysis (IPA). Bar plots display the most significantly enriched categories associated with physiological system development and function. The dashed vertical line indicates the statistical significance threshold (−log10 P = 1.3). (**C**) Disease and function enrichment analysis of differentially expressed proteins using IPA. The most significantly represented disease-associated categories are shown. The dashed vertical line indicates the statistical significance threshold (−log10 P = 1.3). (**D**) Heatmap summarizing significantly altered canonical signaling pathways identified by IPA across the indicated experimental comparisons. Colors correspond to predicted activation states based on IPA activation z-scores, ranging from inhibited (blue) to activated (orange). (**E**) Heatmap of significantly altered metabolic pathways predicted by IPA. Color intensity reflects pathway activation z-score, with blue indicating predicted inhibition and orange indicating predicted activation. NIR, nonirradiated; IR, UVB-irradiated.

**Supplementary Figure S3.**
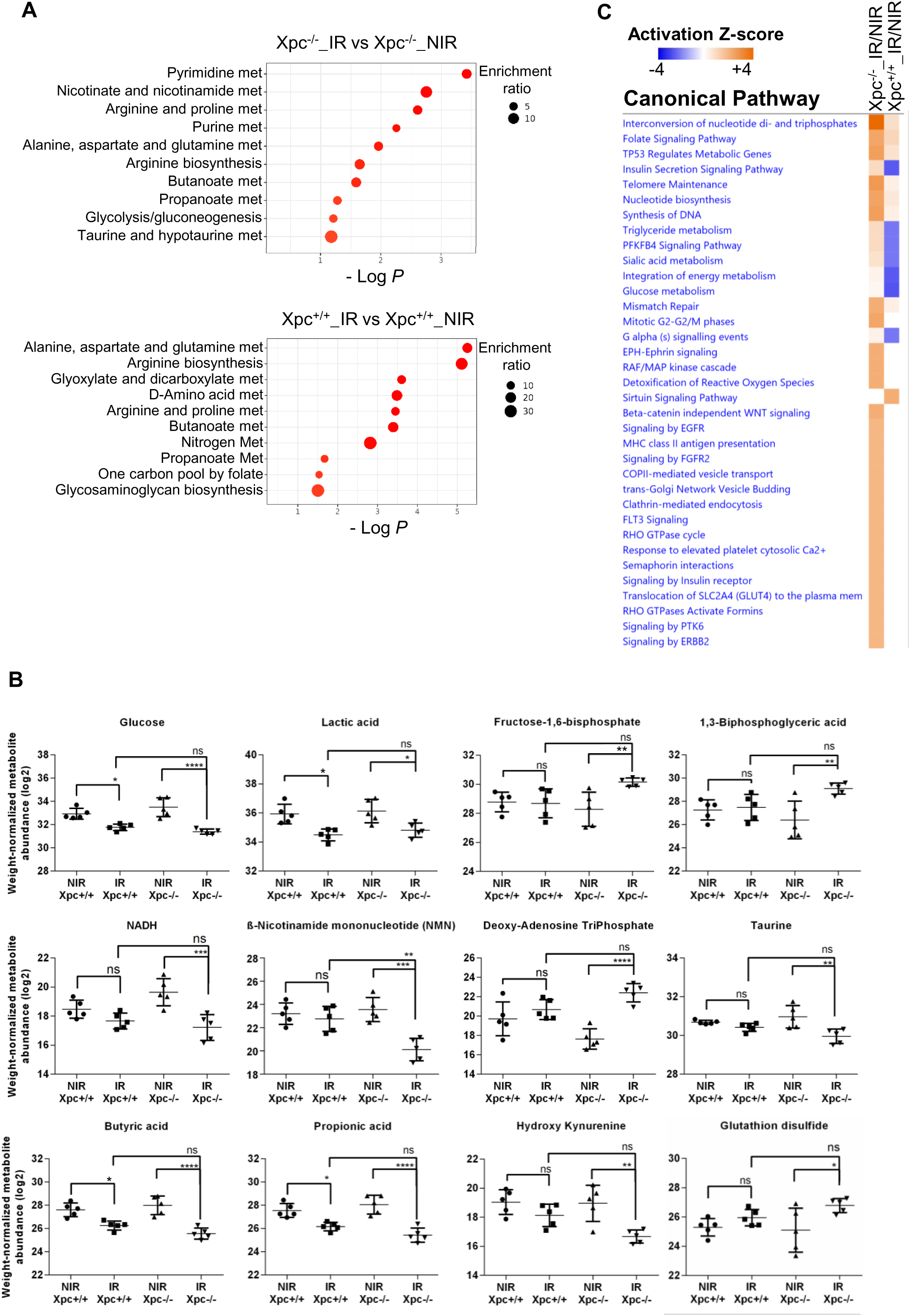
Metabolomic analyses reveal redox imbalance and metabolic adaptation in UVB-irradiated Xpc-deficient skin. (**A**) Metabolic pathway enrichment analysis derived from semi-targeted metabolomic profiling. Bubble plots illustrate significantly enriched metabolic pathways (y-axis) identified in irradiated versus non-irradiated *Xpc*^−/−^ skin (top) and irradiated versus non-irradiated *Xpc*^+/+^ skin (bottom). The x-axis represents pathway enrichment significance (−log10 P value), whereas bubble size reflects pathway enrichment ratio. **(B)** Relative abundance of the indicated metabolites across the four experimental conditions. Metabolite abundances were derived from peak areas normalized to the tissue weight and were subsequently log_2_-transformed. Data are presented as mean ± SD (n = 5 ear skin samples per condition). Statistical significance was assessed by two-way ANOVA followed by Šídák’s multiple comparisons test (* *P* < 0.05, \*\**P* < 0.01, *** *P* < 0.001, **** *P* < 0.0001). (**C**) Heatmap summarizing canonical pathways significantly altered according to metabolomic datasets analyzed using Ingenuity Pathway Analysis (IPA). Pathway activity is represented by IPA activation z-scores, ranging from inhibited (blue) to activated (orange). Comparisons include irradiated versus non-irradiated *Xpc*^−/−^ mice and irradiated versus non-irradiated *Xpc*^+/+^ mice. NIR, nonirradiated; IR, UVB-irradiated.

**Supplementary Figure S4.**
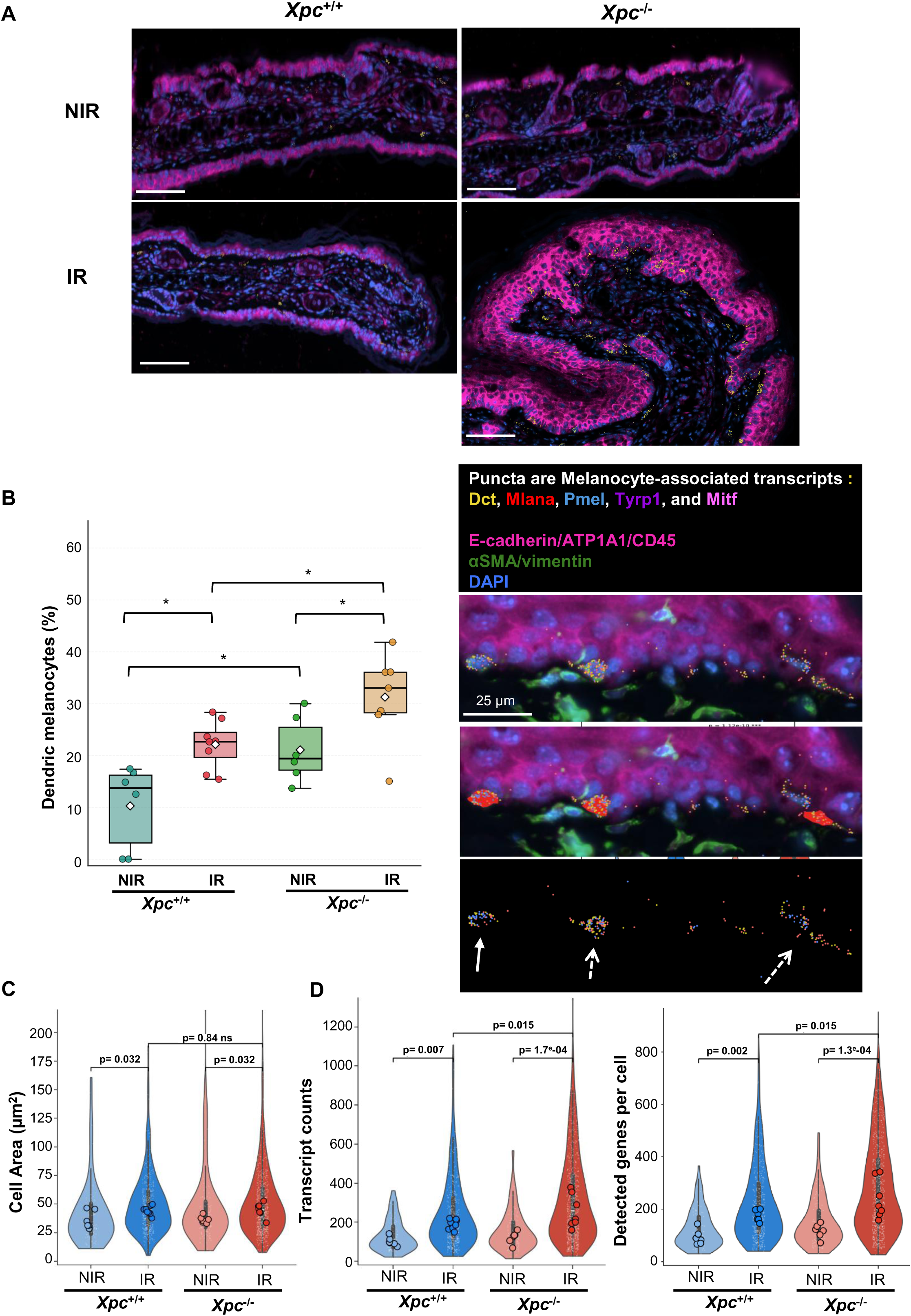
XPC deficiency promotes melanocyte redistribution, dendricity and transcriptional activation following UVB irradiation. **(A)** Representative high-resolution Xenium multimodal image of non-irradiated and irradiated *Xpc*^+/+^ and *Xpc*^−/−^ ear skin. Multimodal staining includes DAPI (blue) and E-cadherin/ATP1A1/CD45 (magenta), together with melanocyte-specific transcripts (*Pmel*, *Mlana*, and *Dct*; yellow puncta), illustrating the spatial distribution of melanocytes across the four experimental conditions. **B)** Boxplots (left) present quantification of dendritic melanocytes across the four experimental conditions. Dendritic melanocytes were manually quantified from high-resolution Xenium multimodal images. Statistical significance was assessed using the Mann–Whitney U test. In the representative image (right), melanocyte-associated transcripts are displayed as *Dct* (yellow), *Mlana* (red), *Pmel* (blue), *Tyrp1* (violet), and *Mitf* (pink), overlaid with E-cadherin/ATP1A1/CD45 (magenta), αSMA/vimentin (green), and DAPI (blue). Dashed arrows indicate dendritic melanocytes, whereas solid arrows indicate non-dendritic melanocytes. **C)** Single-cell quantification of melanocyte surface area (µm^2^) across the four experimental conditions. Statistical significance was assessed using two-way ANOVA. **(D)** Single-cell quantification of total transcript counts per melanocyte and the number of detected genes per melanocyte across the four experimental conditions. Statistical significance was assessed using two-way ANOVA. NIR, nonirradiated; IR, UVB-irradiated.

**Supplementary Figure S5.**
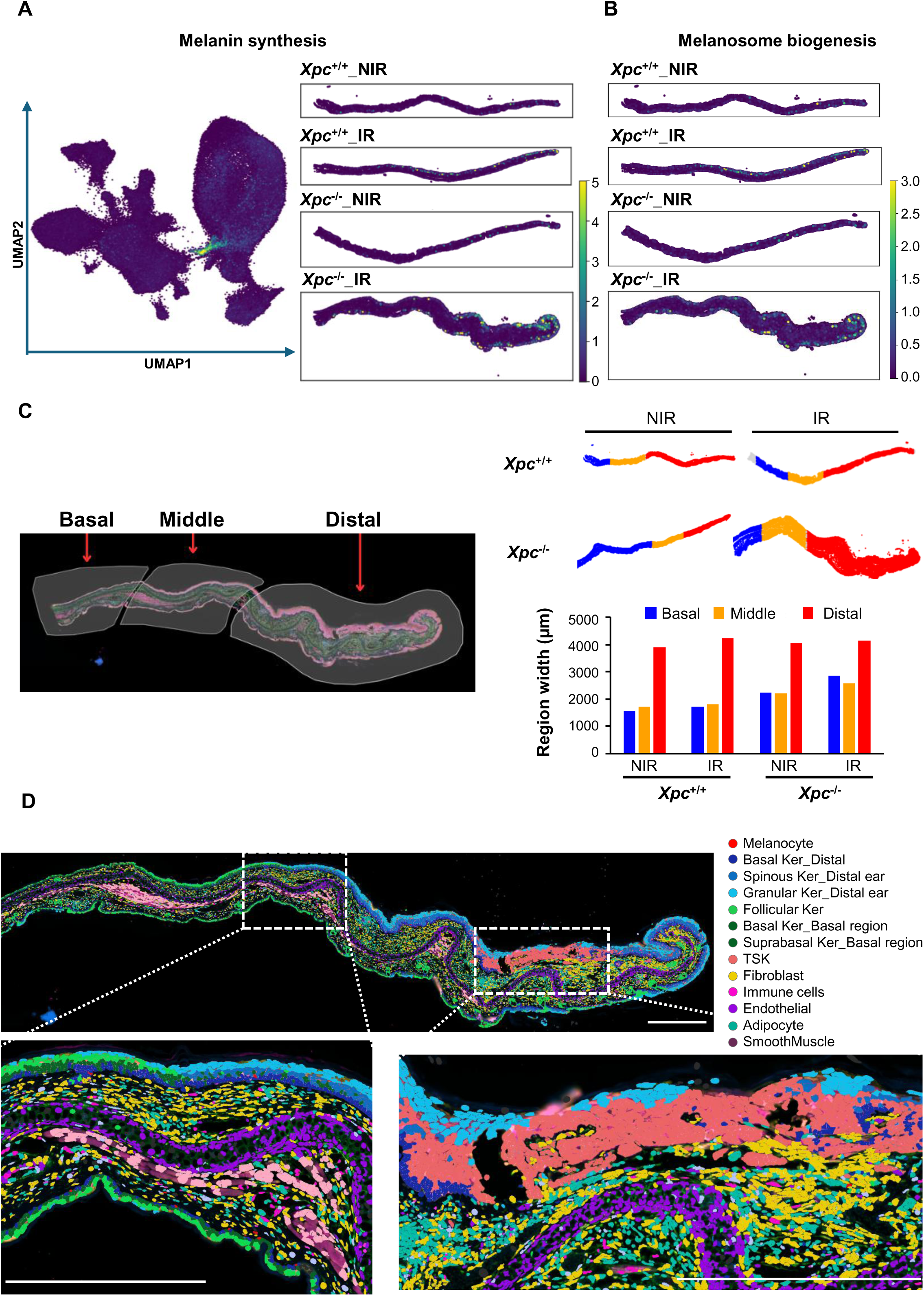
Regional organization of pigmentation-associated programs and anatomical definition of the mouse ear for spatial transcriptomic analyses. (**A–B**) Spatial gene-signature maps generated from Xenium transcriptomic data showing regional enrichment of melanin synthesis (**A**) and melanosome biogenesis (**B**) signatures across the four experimental conditions. Signature scores were calculated from predefined gene sets and projected onto tissue coordinates. Both pigmentation-associated signatures showed pronounced enrichment toward the distal region of irradiated Xpc^−/−^ ears. (**C**) Representative high-resolution Xenium multimodal image of an irradiated Xpc^−/−^ ear illustrating the anatomical subdivision of the tissue into basal, middle, and distal regions used for subsequent regional spatial transcriptomic analyses (left). Spatial maps (top right) illustrate the corresponding basal (blue), middle (orange), and distal (red) regions across the four experimental conditions. Quantification of regional dimensions across experimental groups is shown below. Comparable dimensions were observed across conditions, supporting consistent anatomical subdivision for regional analyses. (**D**) Xenium spatial transcriptomic map showing the distribution of annotated cell types across an irradiated Xpc^−/−^ ear section. Cells are color-coded according to their assigned cell type. Enlarged views show the cellular organization of representative proximal and distal regions. Scale bars, 500 µm. NIR, nonirradiated; IR, UVB-irradiated.

**Figure S6.**
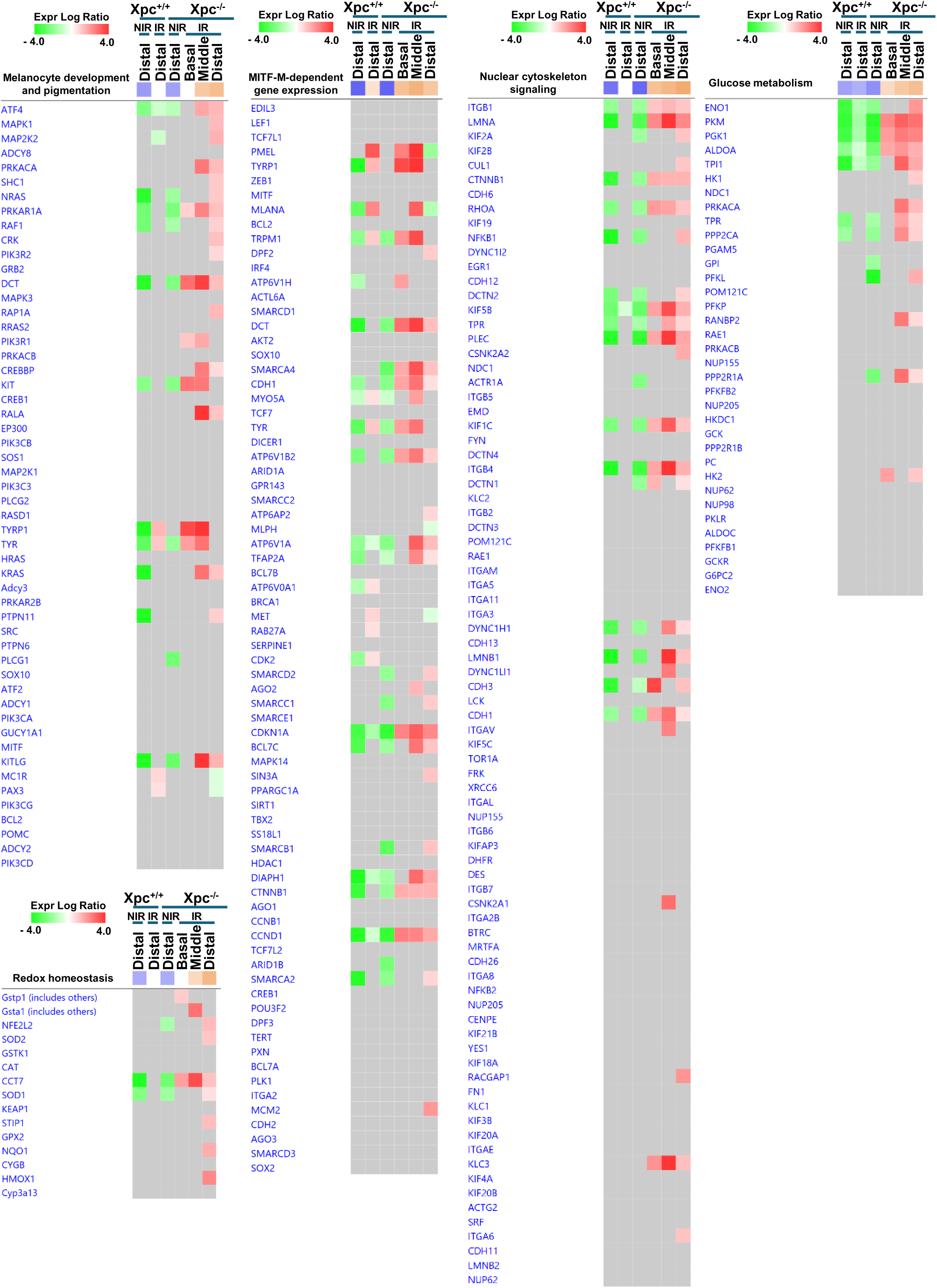
Transcriptional reprogramming of melanocytes in UVB-irradiated *Xpc*-deficient skin. Heatmaps showing representative genes contributing to the major biological pathways identified by Ingenuity Pathway Analysis (IPA) in melanocytes across the four experimental conditions and anatomical regions. Pathways include melanocyte development and pigmentation signaling, MITF-dependent gene expression, cytoskeletal signaling, glucose metabolism, and redox homeostasis. Heatmap colors indicate normalized expression log ratios (green, decreased; red, increased), highlighting the coordinated transcriptional reprogramming of melanocytes in UVB-irradiated *Xpc*^−/−^ skin. NIR, nonirradiated; IR, UVB-irradiated.

**Supplementary Figure S7.**
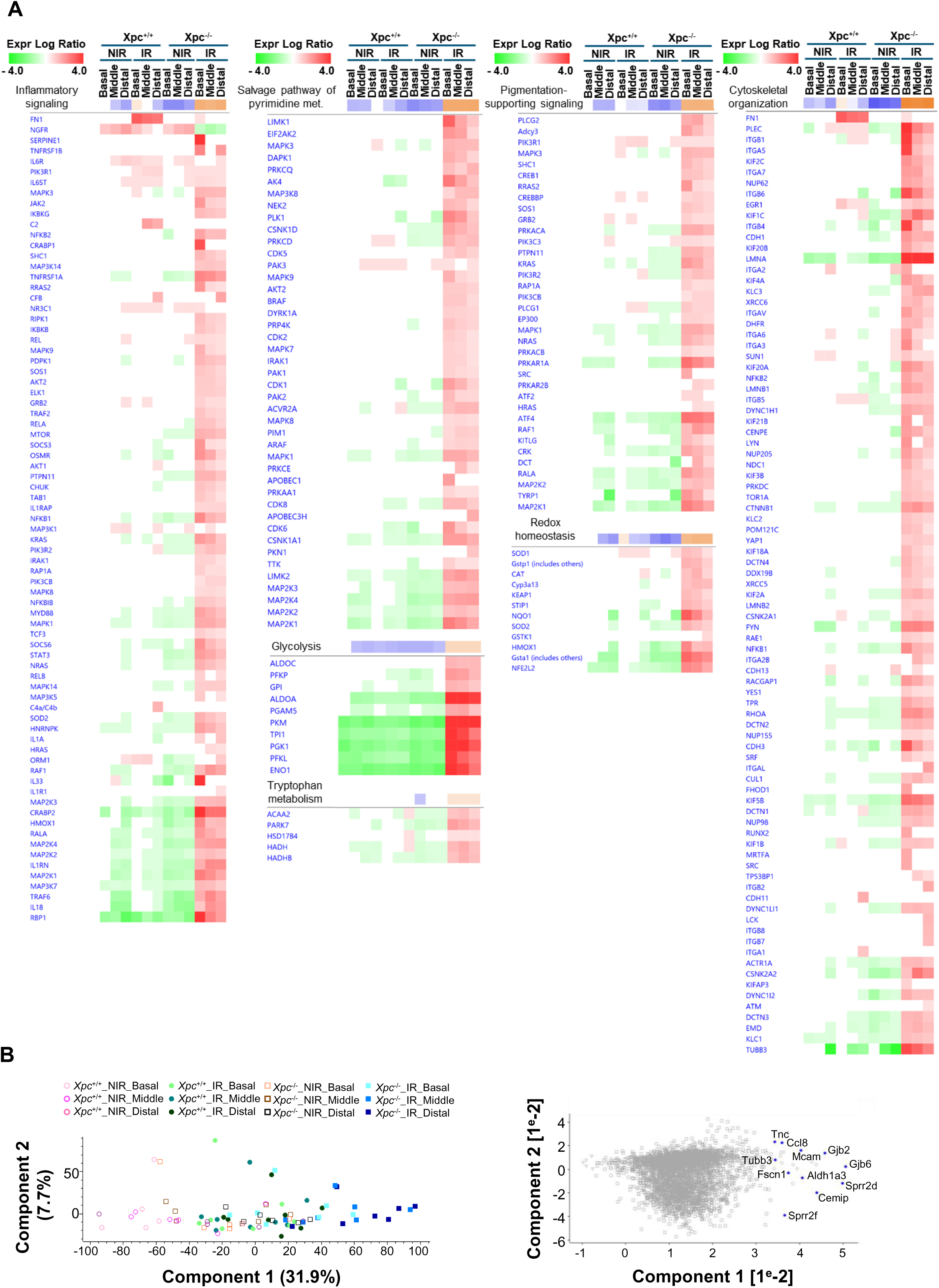
Transcriptional signatures underlying the stress-associated keratinocyte state in UVB-irradiated *Xpc*-deficient skin. (**A**) Heatmaps showing representative genes contributing to major biological pathways identified by Ingenuity Pathway Analysis (IPA) in keratinocytes across the four experimental conditions and anatomical regions. Pathways include inflammatory signaling, pyrimidine salvage, glycolysis, tryptophan metabolism, pigmentation-supporting signaling, redox homeostasis, and cytoskeletal organization. Heatmap colors indicate normalized expression log ratios (green, decreased; red, increased), highlighting coordinated transcriptional alterations in UVB-irradiated *Xpc*−/− keratinocytes. (**B**) Principal component analysis (PCA) of keratinocyte transcriptomic profiles across experimental conditions and anatomical regions (left). UVB-irradiated *Xpc*^−/−^ keratinocytes show distinct transcriptional separation from the other experimental groups, particularly within the middle and distal ear regions. PCA loading analysis (right) identifies genes contributing strongly to this separation, including *Gjb2, Gjb6, Sprr2d, Sprr2f, Cemip, Fscn1, Mcam, Tnc,* Ccl8, and *Aldh1a3*, consistent with a stress-associated epidermal program involving inflammatory signaling, tissue remodeling, and cytoskeletal reorganization. NIR, nonirradiated; IR, UVB-irradiated.

**Supplementary Figure S8.**
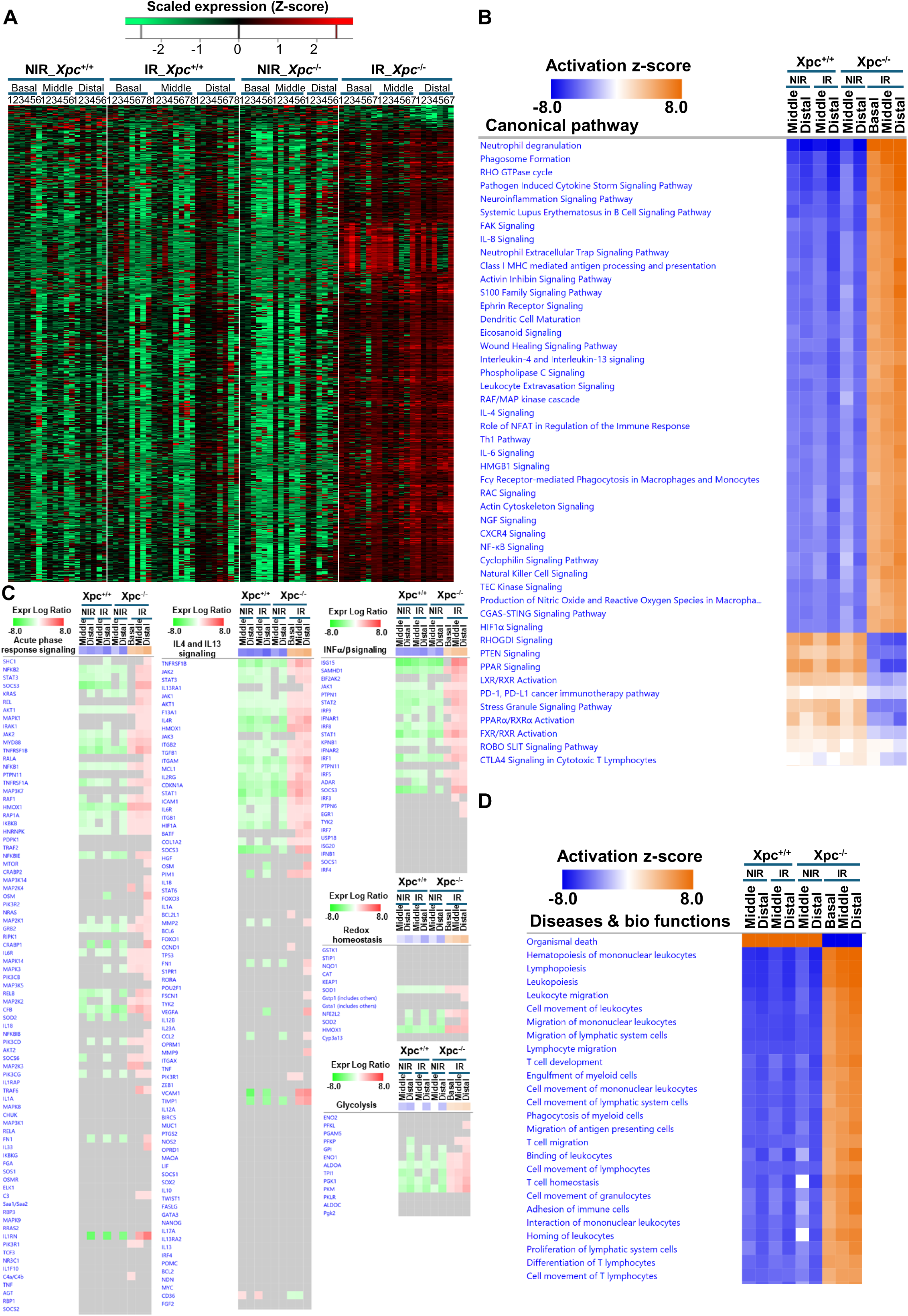
Transcriptional and functional reprogramming of immune cells in UVB-irradiated *Xpc*-deficient skin. All analyses were performed using Xenium spatial transcriptomic data from four experimental groups: *Xpc*^+/+^_NIR (n = 6), *Xpc*^+/+^_IR (n = 8), *Xpc*^−/−^_NIR (n = 6), and *Xpc*^−/−^_IR (n = 7), where n represents the number of mice. **(A)** Heatmap showing gene expression profiles of immune cells across the four experimental conditions and three anatomical ear regions (basal, middle, and distal). Each column represents the aggregated transcript expression profile of immune cells from one mouse within a defined anatomical region. The green-to-red color scale represents normalized, Z-score-transformed transcript abundance, with green indicating lower expression and red indicating higher expression. (**B**) Ingenuity Pathway Analysis (IPA) of canonical pathways associated with differentially expressed genes in immune cells. Orange and blue indicate predicted pathway activation and inhibition, respectively, based on IPA activation z-scores. UVB-irradiated *Xpc*^−/−^ immune cells display coordinated alterations in pathways associated with inflammatory and cytokine signaling, innate and adaptive immune responses, phagocytosis, leukocyte trafficking, and cellular stress responses. (**C**) Heatmaps showing representative genes contributing to selected immune and metabolic pathways identified by IPA, including acute-phase response signaling, IL-4/IL-13 signaling, IFN-α/β signaling, redox homeostasis, and glycolysis. Heatmap colors indicate normalized expression log ratios (green, decreased; red, increased) across anatomical regions and experimental conditions. (**D**) IPA Diseases and Bio Functions analysis of differentially expressed genes in immune cells. Heatmap shows predicted functional changes based on IPA activation z-scores (orange, predicted activation; blue, predicted inhibition). NIR, nonirradiated; IR, UVB-irradiated.

**Figure S9.**
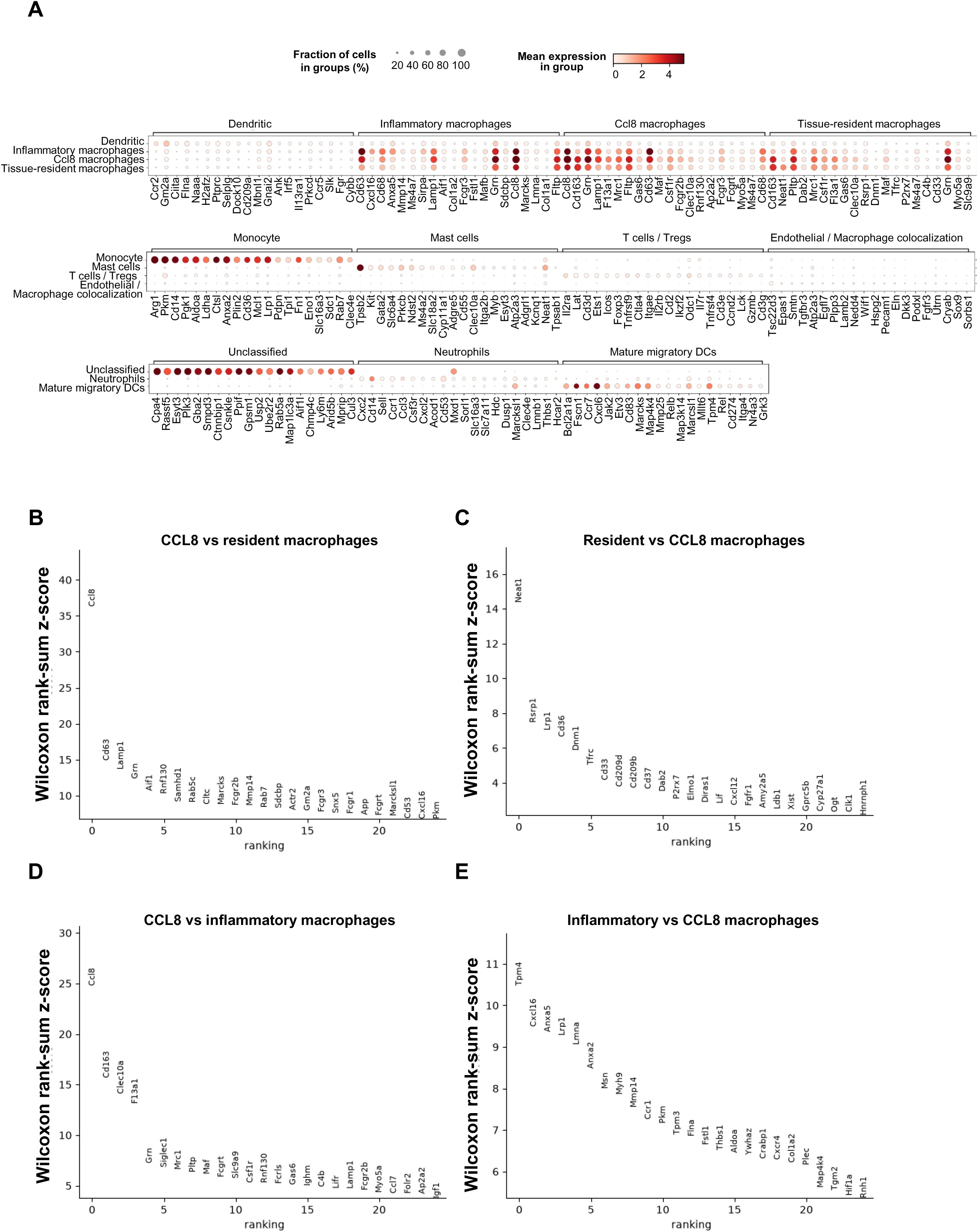
Transcriptional characterization of distinct macrophage states identified by Xenium spatial transcriptomics. **(A)** Dot plot showing expression of selected marker genes across immune-cell populations identified following reclustering of immune cells from the Xenium dataset. Dot size represents the fraction of cells expressing each gene, and color intensity indicates mean expression level. **(B–E)** Differentially expressed genes between the indicated macrophage populations were ranked using the Wilcoxon rank-sum test implemented in Scanpy. The y-axis represents the Wilcoxon test z-score, with higher values indicating stronger evidence for differential expression in the indicated direction. Differentially expressed genes were considered significant at Benjamini–Hochberg-adjusted *P* < 0.05. **(B)** Top differentially expressed genes enriched in CCL8+ macrophages compared with tissue-resident macrophages. CCL8+ macrophages showed increased expression of *Ccl8, Cd63, Lamp1, Grn, Samhd1, Fcgr2b, Mertk, Rab7, Gm2a, Fcgr3, Fcgr1, Fcgrt, Cd53,* and *Cxcl16*, consistent with enhanced chemokine signaling, Fc-receptor function, and endolysosomal/phagocytic activity. **(C)** Top differentially expressed genes enriched in tissue-resident macrophages compared with CCL8+ macrophages, including *Lrp1, Cd36, Cd209d, Cd209b, Dab2, P2rx7, Elmo1, Cxcl12,* and *Cyp27a1*, supporting a transcriptionally distinct resident macrophage state. **(D)** Top differentially expressed genes enriched in CCL8+ macrophages compared with inflammatory macrophages. CCL8+ macrophages showed increased expression of *Ccl8, Cd163, Clec10a, F13a1, Grn, Siglec1, Mrc1, Maf, Csf1r, Fcgr3, Igf1,* and *Folr2*, consistent with a resident/scavenging and tissue-remodeling macrophage phenotype associated with strong *Ccl8* expression. **(E)** Top differentially expressed genes enriched in inflammatory macrophages compared with CCL8+ macrophages. Inflammatory macrophages displayed increased expression of genes associated with cellular stress, glycolytic metabolism, cytoskeletal remodeling, migration, and extracellular-matrix remodeling, including *Cxcl16, Anxa2, Msn, Myh9, Mmp14, Pkm, Flna, Thbs1, Aldoa, Cxcr4, Tgm2,* and *Hif1a*.

**Figure S10.**
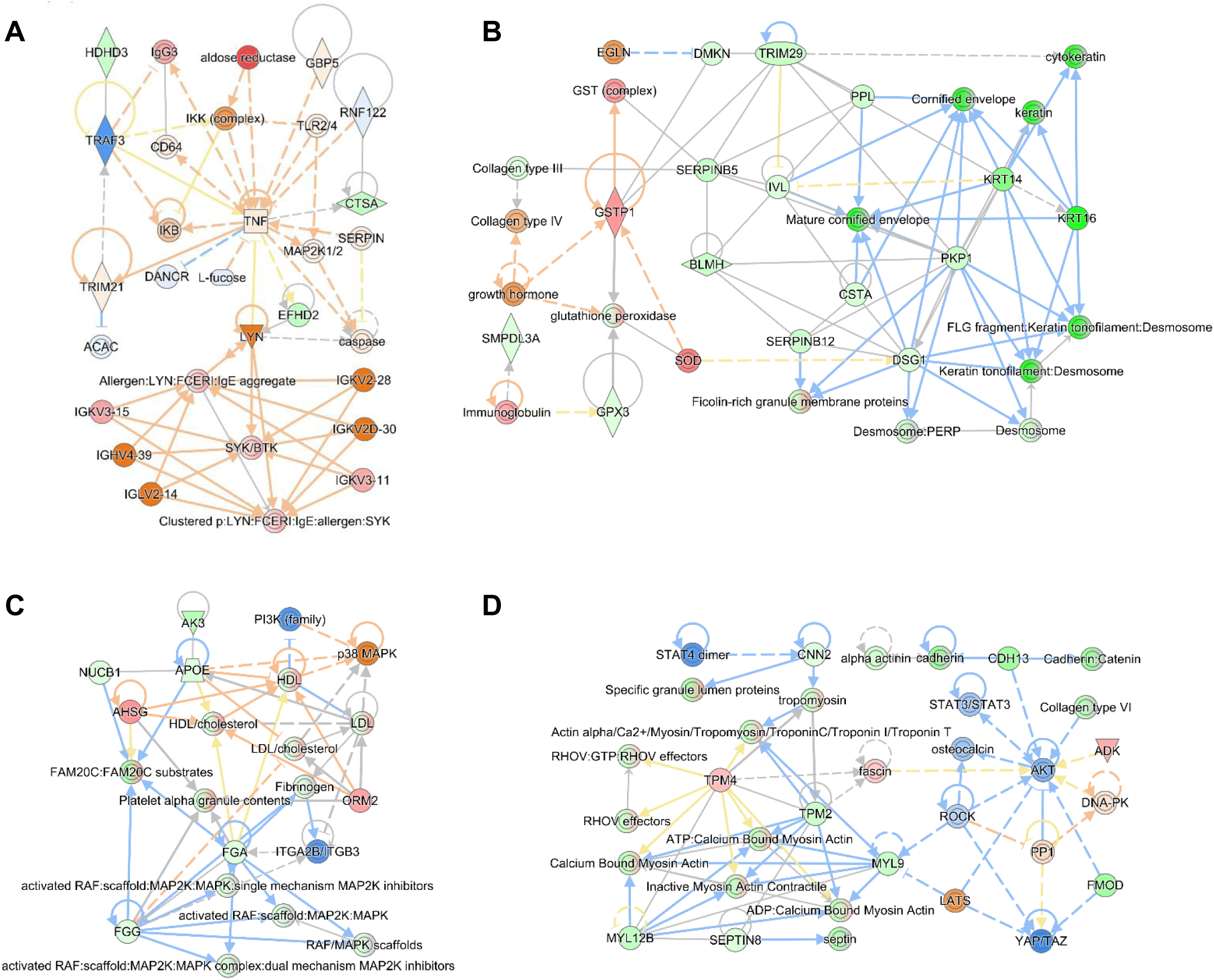
Proteomic profiling reveals conserved inflammatory, keratinization and epidermal differentiation, metabolic rewiring, and cytoskeletal remodeling in human XP-C hyperpigmented skin. **(A–D)** Representative mechanistic networks generated by IPA illustrating the major biological processes altered in hyperpigmented XP-C skin. Node colors indicate measured protein abundance (red, increased; green, decreased), whereas pathway nodes are colored according to their predicted activation state (orange, activated; blue, inhibited). Solid and dashed lines indicate direct and indirect molecular interactions, respectively. Networks illustrate coordinated remodeling of (**A**) inflammatory and immune signaling centered on TNF and SRC-family kinase pathways, (**B**) epidermal differentiation and keratinization, (**C**) lipid metabolism and MAP kinase signaling, and (**D**) actomyosin cytoskeletal organization, cell-matrix adhesion and mechanotransduction pathways.

## Supplementary Materials and Methods

### Animals and UVB irradiation

XPC-deficient mice were generated from *Xpc* heterozygous (*Xpc*^+/−^) breeders originally described by Cheo et al. ^41^ and generously provided by Errol C. Friedberg (The University of Texas Southwestern Medical Center, Dallas, TX, USA). The colony has been maintained in our animal facility for more than 10 years on a C57BL/6J genetic background ^42^. The wild-type (*Xpc*^+/+^) and knockout (*Xpc*^−/−^) littermates were generated for all experiments by breeding *Xpc*^+/−^ mice. Genotypes were confirmed by PCR analysis of genomic DNA obtained from tail biopsies using primers described previously ^42^. A total of 70 female mice were included in three independent biological cohorts. Each experiment comprised four experimental groups: non-irradiated wild-type (*Xpc*^+/+^-NIR), irradiated wild-type (*Xpc*^+/+^-IR), non-irradiated knockout (*Xpc*^−/−^-NIR), and irradiated knockout (*Xpc*^−/−^-IR), with 5–8 mice per group depending on the downstream analyses performed. All animals were housed in groups of five mice per cage under specific pathogen-free conditions at the University of Bordeaux animal facility, under controlled temperature, humidity and a 12-hour light/dark cycle, with ad libitum access to food and water.

Eight- to ten-week-old female *Xpc*^+/+^ and *Xpc*^−/−^ mice were subjected to a chronic UVB irradiation protocol. For each irradiation session, mice were placed individually in custom cages to ensure homogeneous UVB exposure. Irradiated mice were pooled and subjected to UVB exposure with the investigators blinded to genotype; non-irradiated mice were handled similarly under genotype-blinded conditions. Animals received UVB doses of 75 mJ/cm^2^ on days 0, 2 and 5, followed by 100 mJ/cm^2^ on days 7, 9 and 12. Pigmentation was monitored throughout the experiment by serial photography of the ears, dorsal skin and tail. Of note, investigators remained blinded to genotype during the initial irradiation period; however, blinding could no longer be ensured after the emergence of the visible hyperpigmentation phenotype. Mice were euthanized on day 14 by cervical dislocation, and tissues were immediately harvested and processed for histological, ultrastructural, molecular and multi-omics analyses.

All animal experiments were performed in accordance with institutional guidelines and approved by the University of Bordeaux Animal Care and Use Committee, in compliance with national and European regulations governing the use of laboratory animals.

### Isolation and primary culture of murine melanocytes

Primary melanocytes were isolated from mouse ear skin using a combined mechanical and enzymatic dissociation protocol (**supplementary Fig. S1**). Briefly, freshly dissected ears were rinsed in ice-cold Dulbecco’s phosphate-buffered saline (DPBS) and minced into approximately 1 mm^3^ fragments using sterile scalpels. Tissue fragments were incubated for 90 min at 37°C in DPBS containing 1.25 mg/mL collagenase (Sigma-Aldrich, Cat# C9891), 2mg/mL dispase (Sigma-Aldrich, Cat# P3417) and 1 mM CaCl_2_, with gentle manual agitation every 30 min.

Following enzymatic digestion, tissues were mechanically dissociated by repeated trituration using a 10-mL serological pipette until a homogeneous cell suspension was obtained. The suspension was filtered through a 70-μm cell strainer, washed three times with DPBS, and centrifuged at 100 × g for 10 min. The resulting cell pellet was washed three additional times with DPBS before being resuspended in Medium 254 (Thermo Fisher Scientific) supplemented with Human Melanocyte Growth Supplement (HMGS, Thermo Fisher Scientific) and 1% penicillin–streptomycin, which allowed preferential expansion of primary melanocytes.

The culture medium was completely replaced after 24 h to remove non-adherent cells and cellular debris and was subsequently refreshed every 2–3 days. Cells were maintained at 37°C in a humidified incubator containing 5% CO_2_. For passaging, cells were washed with phosphate-buffered saline (PBS) and detached using 0.05% trypsin–EDTA (Gibco) for 5 min at 37°C. Trypsinization was terminated with complete culture medium, and cells were collected by centrifugation at 100 × g for 5 min before reseeding or use in subsequent experiments. Primary melanocytes between passages 1-3 were used for all experiments. Melanocyte identity was confirmed by immunofluorescence staining for TYR, MITF and c-KIT as shown in **Supplementary Fig. S1**.

### Live-cell imaging of melanosome transfer

Primary murine melanocytes isolated from the ear skin of *Xpc*^+/+^ and *Xpc*^−/−^ mice were seeded and cultured to approximately 60% confluence. Cells were then transduced with MelanoTag-PURO lentivirus (pLJC5-GPR143-mScarlet-3xHA; a gift from David Sabatini; Addgene plasmid # 160369; http://n2t.net/addgene:160369) ^16^ to stably express the GPR143-mScarlet-3xHA fusion protein, enabling fluorescent labeling and tracking of melanosomes. After 24 h, the virus-containing medium was replaced with fresh growth medium, and transduced melanocytes were maintained for an additional 72 h before expansion for subsequent experiments. In parallel, PDVC57 murine keratinocytes ^43^ were stably transduced with a lentiviral vector encoding enhanced green fluorescent protein (EGFP), as previously described ^44^.

To visualize melanosome transfer from melanocytes to recipient keratinocytes, MelanoTag-labeled melanocytes and EGFP-expressing keratinocytes were cocultured at a 1:10 melanocyte-to-keratinocyte ratio in six-well µ-Slide chambers (ibidi). Cocultures were exposed to UVB at 5 mJ/cm^2^ every 48 h for a total of three irradiations to stimulate pigmentation and melanosome transfer. Live-cell imaging was performed immediately after the third UVB exposure, with approximately 15 fields of view acquired per well to monitor the dynamics of fluorescently labeled melanosomes and their transfer from melanocytes to neighboring keratinocytes.

### Metabolomic analyses

Metabolites were extracted from mouse ear tissues (n = 5 mice per group) using the EtOH/HEPES protocol described by Ceschin et al ^45^. Briefly, pre-weighed frozen ears were homogenized using a Tissue Lyzer (QIAGEN) at 30 Hz for 2 × 30 s. Metabolites were extracted with EtOH/1 mM Hepes, pH 7.0 (8:2, v/v), containing the heavy isotope-labeled internal standards succinic acid-2,2,3,3-D_4_, L-aspartic acid-^13^C_4_,^15^N, L-tyrosine-^13^C_9_, and adenosine-2D_14_ 5′-triphosphate. Samples were heated at 80 °C for 3 min and dried in a vacuum concentrator at 65 °C for 30 min. Dried extracts were reconstituted in Milli-Q water at 20 µl/mg of tissue. Insoluble material was removed by centrifugation at 21,000 × g for 1 h at 4 °C, and the resulting supernatants were subjected to ultrafiltration using Nanosep 10K Omega centrifugal devices (Pall) at 14,000 × g for 15 min at 4 °C. Metabolites were separated by either ultra-high-performance ion chromatography (u-HPIC; ICS6000, Thermo Fisher Scientific) or ultra-high-performance liquid chromatography (u-HPLC; Vanquish Flex, Thermo Fisher Scientific), using an AS11-HC column (4 µm, 250 × 2 mm) or an Acclaim RSLC PAII column (2.2 µm, 120 Å, 2.1 × 150 mm), respectively. HPIC separation was performed at a flow rate of 0.38 mL/min using a discontinuous KOH gradient, as previously described ^46^. For HPLC separation, metabolites were resolved of 0.25 mL/min using 50 mM formic acid (pH 2.9, adjusted with ammonia) as mobile phase A and methanol as mobile phase B, according to the gradient described previously ^47^. For both chromatographic methods, metabolites were detected using high-resolution Orbitrap mass spectrometer (Exploris 120, Thermo Fisher Scientific) equipped with an EASY-IC ion source. Data were acquired in negative-ion mode for HPIC and in both negative- and positive-ion modes for UHPLC, with scan-to-scan lock-mass correction. Data acquisition was performed using Xcalibur 4.7 (Thermo Fisher Scientific). Targeted metabolites were quantified from full-scan MS data (m/z 70–1,000) using TraceFinder 5.2 SP3 (Thermo Fisher Scientific). Full-scan spectra were acquired at a resolution of 60,000, with data-dependent MS^2^ spectra acquired at a resolution of 15,000 using higher-energy collisional dissociation (HCD) at a normalized collision energy of 30%. Metabolite identification was based on retention time, accurate mass, natural isotopic distribution, and MS^2^ fragmentation patterns. Peak areas were normalized to the corresponding heavy isotope-labelled internal standards. Because the reconstitution volume was adjusted to the initial tissue weight (20 µL/mg), the resulting values represent tissue weight-normalized metabolite abundances.

Metabolite abundance data were subsequently analyzed using Perseus (v2.1.6.0) ^48^. Tissue weight-normalized metabolite abundances were log_2_-transformed before statistical analysis. Metabolites with missing values in more than 50% of all samples were excluded, and remaining missing values were imputed from a normal distribution using a width (standard deviation) of 0.3 and a downshift of 1.8 (mean adjustment of −1.8 units). Hierarchical clustering was computed using Spearman correlation as the distance measure and average linkage.

Differentially abundant metabolites (DAMs) were identified by pairwise comparison of nonirradiated and irradiated *Xpc*^−/−^ samples and nonirradiated and irradiated *Xpc*^+/+^ samples using two-tailed Student’s *t*-tests with permutation-based FDR correction (250 randomizations; FDR = 0.05). DAMs were subsequently subjected to metabolic pathway enrichment analysis using MetaboAnalyst (https://www.metaboanalyst.ca) and the KEGG metabolic pathway library (November 2025 release). DAMs were also subjected to canonical pathway and functional enrichment analyses using Ingenuity Pathway Analysis (IPA; QIAGEN).

For integrated multi-omics analysis, proteomic, metabolomic, and kinome datasets were analyzed using the Joint Pathway Analysis module of MetaboAnalyst. Differentially expressed proteins (DEPs), differentially abundant metabolites (DAMs), and predicted upstream kinases were jointly mapped to KEGG pathways and analyzed using pathway enrichment and topology-based analysis.

### Proteomic analyses

#### 1-Sample Preparation

- **Primary Human Melanocyte CRISPR-Cas9 Model.** Primary human melanocytes derived from five independent donors were subjected to a CRISPR-Cas9-mediated knockout, generating five matched biological pairs of control (Ctrl) and XPC-deficient (XPC-KO) melanocytes, as already described ^18^. Cells were lysed in RIPA buffer (50 mM Tris-HCl, pH 8.0; 150 mM NaCl; 0.1% SDS; 1% NP-40; 0.5% sodium deoxycholate; protease inhibitor cocktail, Sigma-Aldrich) using a Precellys homogenizer with CK14 beads (Bertin, Ozyme, France). Lysates were centrifuged at 12,500 rpm for 20 min at 4°C, and protein concentrations were determined using a BCA assay (Pierce, Thermo Fisher Scientific).
- **XPC patient skin biopsies.** Biopsy samples were obtained from three XP-C patients (N = 3, supplementary table 1). For each patient, biopsies were collected from hypopigmented and hyperpigmented skin lesions. Tissue samples were processed for protein extraction and solubilized in Laemmli bufferand subjected to SDS-PAGE.
- **Mouse ear skin.** Ear skin samples were collected from the experimental mouse cohorts described above. For the proteomic analysis, five biological replicates per experimental group were analyzed (n = 5 mice per group). Ear tissues were solubilized in Laemmli buffer and subjected to SDS-PAGE.

#### 2-In-gel proteolytic Digestion

Equal amounts of protein were seprataed by SDS-PAGE on 10% polyacrylamide gels and stained with colloidal Coomassie blue. Individual gel lanes were excised and cut into approximately 1 × 1 mm gel pieces. The pieces were destained in 25 mM NH4HCO3 containing 50% acetonitrile (CAN), rinsed twice with ultrapure water and dehydrated in 100% ACN for 10 min. After removal of the supernatant and air-drying at room temperature, gel pieces were covered with sequencing-grade trypsin solution (10 ng/µL in 50 mM NH4HCO3), allowed to rehydrate for 10 min at 4°C, and incubated overnight at 37°C. Peptides were first extracted by incubating the gel pieces in 50 mM NH4HCO3 for 15 min under gentle agitation. The supernatant was collected, and two additional extractions were performed using H_2_O/ACN/formic acid (47.5:47.5:5, v/v/v) for 15 min each. The combined peptide extracts were pooled, dried in a vacuum centrifuge and resuspended in 0.1% formic acid before LC–MS/MS analysis.

#### 3-Nano-LC-MS/MS Analysis

Peptide mixtures were analyzed using an Ultimate 3000 RSLC nanoUHPLC system (Thermo Fisher Scientific / Dionex, Amsterdam, The Netherlands) coupled to an Orbitrap Fusion Lumos Tribrid mass spectrometer (Thermo Fisher Scientific, San Jose, CA). All samples were initially loaded onto a PepMap C18 trap column (300 µm × 5 mm, LC Packings) at a flow rate of 10 µL/min and desalted for 3 min.

##### Data-dependent acquisition of human melanocyte and patient biopsy samples

Peptides were separated on a 50-cm EASY-Spray analytical column (75 µm internal diameter, 2 µm particles) using a linear gradient from 4% to 40% solvent B (0.1% formic acid in 80% CAN) over 48 min at a flow rate of 300 nL/min. The mass spectrometer was operated in positive-ion data-dependent acquisition (DDA) mode with a spray voltage of 2.0 kV. Full MS1 spectra were acquired over an m/z range of 375–1,500) in the Orbitrap at a resolution of 120,000 (at m/z 200), with an automatic gain control (AGC) target of 4 × 10^^5^ and a maximum injection time of 50 ms. MS/MS acquisition was performed using a top-speed method with a 3-s cycle time. Precursor ions with charge states 2–7 were isolated using a 1.6-Th window and fragmented by higher-energy collisional dissociation (HCD) at a normalized collision energy (NCE) of 28%. MS/MS spectra were acquired in the ion trap using dynamic injection times, with a dynamic exclusion window of 60 s.

##### Data-independent acquisition of mouse ear skin samples

Peptides were separated on a 75 µm × 60 cm C18 analytical column (IonOpticks, Australia). Chromatographic separation was performed using a linear gradient of solvent B (0.1% formic acid in 80% ACN) from 4% to 25% over 87 min, followed by an increase from 25% to 40% over 30 min, at a constant flow rate of 200 nL/min. The mass spectrometer was operated in positive-ion data-independent acquisition (DIA) mode under Xcalibur 4.4 control. Each acquisition cycle consisted of a full MS1 scan (resolution, 60,000; AGC target, 4 × 10^^5^; injection time 50 ms; scan range 350–1400 m/z) followed by 34 DIA MS/MS scans (resolution 30,000; AGC target 5 × 10^^5^; maximum injection time, 50 ms; scan range, m/z 350–1,400), followed by 34 DIA MS/MS scans (resolution, 30,000; AGC target, 5 × 10^5^; maximum injection time, 54 ms) using variable-width isolation windows covering the m/z 400– 900 range. HCD fragmentation was performed at a normalized collision energy of 25%.

#### 4-Proteomic data processing and statistical analysis

##### Human proteome processing (DDA data)

Raw mass spectrometry files generated from primary human melanocytes and XP-C patient biopsies were processed using Proteome Discoverer 2.3 (Thermo Fisher Scientific). Peptide identification was performed using the MS Amanda 2.0, Sequest HT, and Mascot 2.5 search engines against the UniProt *Homo sapiens* reference database (74,489 entries; May 2019). Searches were performed with trypsin specificity, allowing up to two missed cleavages. Precursor and fragment mass tolerances were set to 10 ppm and 0.6 Da, respectively. Carbamidomethylation of cysteine residues (+57.021 Da) was specified as a fixed modification, whereas methionine oxidation (+15.995 Da) and lysine acetylation (+42.011 Da) were included as variable modifications. Peptide-spectrum matches were validated using Percolator with a false discovery rate (FDR) of <1% at the peptide level. Label-free quantification (LFQ) was performed using the Minora Feature Detector node based on precursor-ion intensities. Inter-run variation was normalized to total peptide abundance, and protein abundances were calculated from the summed intensities of the corresponding peptides. The mass spectrometry proteomics data have been deposited with the ProteomeXchange Consortium via the PRIDE repository under dataset identifier PXD042713 ^18^.

Protein abundance data were subsequently analyzed using Perseus (v2.1.6.0). Non-master proteins, proteins identified with fewer than two unique peptides, and proteins annotated as contaminants were excluded before statistical analysis. Proteins with missing values in more than 50% of all samples were excluded. Remaining missing values were imputed from a normal distribution using a width of 0.3 and a downshift of 1.8. Pairwise comparisons were performed between the relevant experimental conditions (Ctrl versus XPC-KO melanocytes and hypopigmented versus hyperpigmented XP-C skin). Statistical significance was assessed using two-tailed Student’s *t*-tests, with permutation-based FDR correction (250 randomizations; FDR = 0.05). Hierarchical clustering was computed using Spearman correlation as the distance measure and average linkage. Differentially expressed proteins (DEPs; FDR < 0.05) were subjected to canonical pathway and functional enrichment analyses using IPA (QIAGEN).

##### Mouse proteome processing (DIA data)

DIA data generated from mouse ear tissues were processed using the CHIMERYS search engine within Proteome Discoverer 3.2 (Thermo Fisher Scientific) against the *Mus musculus* reference proteome database (June 2024 release; 21,736 entries). Peptides with molecular masses below 350 Da or above 5,000 Da were excluded from the database search. The precursor mass tolerance was set to 10 ppm, and up to two missed tryptic cleavages were allowed. Carbamidomethylation of cysteine residues (+57.021 Da) was specified as a fixed modification and methionine oxidation (+15.995 Da) as a variable modification. Peptide identifications were filtered at an FDR of <1%. Peptide and protein abundances were quantified using the Fragment Ion Quantifier node, and inter-sample variation was normalized based on total peptide abundance. The mass spectrometry proteomics data have been deposited with the ProteomeXchange Consortium ^49^ via the PRIDE repository ^50^ under dataset identifier PXD083977. Protein abundance data were subsequently analyzed using Perseus (v2.1.6.0). Non-master proteins, proteins identified with fewer than two unique peptides, and proteins annotated as contaminants were removed. Proteins with missing values in more than 50% of all samples were excluded. Remaining missing values were imputed from a normal distribution using a width of 0.3 and a downshift of 1.8. Principal component analysis (PCA) was performed in Perseus using the processed protein abundance matrix to assess global similarities and differences among samples. Hierarchical clustering was computed using Spearman correlation as the distance measure and average linkage. Pairwise comparisons between experimental groups were performed using two-tailed Student’s *t*-tests with permutation-based FDR correction (250 randomizations; FDR = 0.05; S0 = 0.1). Differences across all experimental groups were assessed by one-way ANOVA with permutation-based FDR correction (250 randomizations; FDR = 0.05; S0 = 0). Differentially expressed proteins (DEPs; FDR < 0.05) were subsequently subjected to canonical pathway and functional enrichment analyses and molecular interaction network analysis using IPA (QIAGEN).

### Functional kinome profiling and upstream kinase analysis

Ear skin tissues were collected at the experimental endpoint from four groups of C57BL/6J mice (n = 6 mice per group). Tissues were immediately flash-frozen and subsequently lysed in M-PER Mammalian Protein Extraction Reagent (Thermo Fisher Scientific, Cat. #78503) supplemented with Halt Protease and Phosphatase Inhibitor Cocktail (Thermo Fisher Scientific). Homogenates were clarified by centrifugation at 4 °C, and total protein concentrations were determined before kinome analysis.

Functional protein tyrosine kinase activity was profiled using the PamChip microarray platform (PamGene International B.V.,’s-Hertogenbosch, The Netherlands) with Protein Tyrosine Kinase (PTK) arrays containing 144 peptide substrates representing distinct phosphorylation sites. For each array, 5 µg of clarified tissue lysate was incubated in kinase assay buffer containing 100 µM ATP and a fluorescein-labeled anti-phosphotyrosine antibody. Assays were performed according to the manufacturer’s protocol. The reaction mixture was repeatedly pumped through the porous three-dimensional array, allowing active kinases in the tissue lysates to phosphorylate immobilized peptide substrates. Peptide phosphorylation was monitored kinetically by fluorescence imaging at predefined time points.

Fluorescence signals were quantified using BioNavigator software (v6.3; PamGene International B.V.). Image processing included kinetic signal extraction, background correction, and integration of signals acquired at multiple exposure times. Peptides not meeting the predefined quality-control criteria were excluded from subsequent analyses. Differential peptide phosphorylation between experimental groups was assessed using empirical Bayes linear modeling implemented in BioNavigator.

Upstream Kinase Analysis (UKA) was subsequently performed in BioNavigator to predict kinases potentially responsible for the observed phosphorylation profiles. Differential peptide phosphorylation patterns were mapped to curated kinase–substrate relationships derived from databases including OmniPath, PhosphoSitePlus, and Reactome. Predicted upstream kinases were ranked using the BioNavigator scoring framework, which integrates the specificity of kinase– substrate relationships with the statistical significance and consistency of phosphorylation changes. Kinases with a median final score >1.25 were considered significantly modulated upstream kinase candidates. To visualize predicted kinase activity changes in the context of kinase family relationships, significantly modulated upstream kinases were mapped onto the human kinome tree using the web-based CORAL tool (http://phanstiel-lab.med.unc.edu/CORAL) ^51^. Kinase node color indicates the predicted direction of activity change, whereas node size is proportional to the UKA median final score.

### Western blot

#### Protein Extraction

Ear skin samples were homogenized in hypotonic buffer (2.5 mM Tris/HCl, pH 7.5 and 2.5 mM MgCl2), supplemented with protease and phosphatase inhibitors (Halt™ Protease and Phosphatase Inhibitor Cocktail, Thermo Fisher Scientific) using a Precellys® Evolution homogenizer. Homogenates were centrifuged at 13 000 rpm for 5 min at 4°C, and the supernatants were collected. A 10x RIPA-like buffer (500 mM Tris-HCl pH 7.4, 1.5 M NaCl, 10% Triton X-100, 1% SDS, and 5% sodium deoxycholate) was added to obtain a 1x final concentration. Samples were incubated on ice for 20 min and centrifuged at 13000 rpm for 20 min at 4°C. The resulting supernatants were collected, and protein concentrations were determined using the BCA protein assay kit (Thermo Fisher Scientific).

#### SDS-PAGE and Transfer

Equal amounts of protein (30 µg) were mixed with 6x Laemmli sample buffer, heated at 95°C for 5 min, and separated on 4–20% Mini-PROTEAN® TGX™ Stain-Free Precast Gels (Bio-Rad). Electrophoresis was performed at 55–120 V until appropriate separation was achieved. Proteins were transferred onto nitrocellulose membranes (Amersham™ Protran®, Merck) using an eBlot™ L1 Protein Transfer System (GenScript).

#### Blocking and Antibody Incubation

Membranes were blocked with 2% ECL Prime Blocking Reagent (Cytiva) for 1 hour at room temperature and subsequently incubated overnight at 4 °C with primary antibodies. The following day, membranes were washed three times for 10 min with TBST and incubated with the appropriate horseradish peroxidase (HRP)-conjugated secondary antibodies for 1 h at room temperature. Details of the primary and secondary antibodies used, including working dilutions, are provided in **the table below**. After additional washes with TBST, protein bands were detected using enhanced chemiluminescence reagent (Biorad) and visualized using an iBright Imaging System (Thermo Fisher Scientific).

### Antibodies used in this study

| Experiment | Antibody-Host Species | Dilution | Supplier |
|---|---|---|---|
| Western Blot | Anti-MITF (R) | 1/1000 | Abcam ab20663 |
| Western Blot | Anti-TRP-1 (R) | 1/1000 | Abcam ab178676 |
| Western Blot | Anti-Tyrosinase (M) | 1/1000 | Bio-techne NBP2-67233 |
| Western Blot | Anti-Rab9A (R) | 1/1000 | Cell Signaling D52G8 |
| Western Blot | Anti-Melan-A (M) | 1/500 | Dako A103 |
| Western Blot | Anti-Rab27a (M) | 1/200 | Santa Cruz sc-74586 |
| Western Blot | Secondary Horse anti-Mouse IgG (H+L) peroxidase | 1/5000 | Vector laboratories PI-2000 |
| Western Blot | Secondary Goat anti-Rabbit IgG (H+L) peroxidase | 1/10000 | Vector laboratories PI-1000-1 |
| Western Blot | Anti-SOX10 | 1/1000 | Abcam ab155279 |
| Western Blot | Anti-β-Actin (ACTB) | 1/5000 | Sigma-Aldrich A2228 |
| Immunofluorescence | Anti-Keratin 10 (R) | 1/200 | Abcam ab76318 |
| Immunofluorescence | Anti-Keratin 14 (M) | 1/100 | Abcam ab7800 |
| Immunofluorescence | Anti-CPD (M) | 1/200 | Cosmo Bio Clone TDM-2 |
| Immunofluorescence | Anti-Melan-A (M) | 1/100 | Dako A103 |
| Immunofluorescence | Anti-Rab27a (M) | 1/100 | Santa Cruz sc-74586 |
| Immunofluorescence | Anti-TRP-1 (R) | 1/1000 | Abcam ab178676 |
| Immunofluorescence | Anti-E-Cadherin (R) | 1/200 | Protein-tech 20874-1-AP |
| Immunofluorescence | Secondary Donkey<br>anti-Mouse IgG (H+L) Antibody,<br>Alexa Fluor™ 568 | 1/500 | Invitrogen A10037 |
| Immunofluorescence | Anti-MITF (R) | 1/200 | Abcam ab20663 |
| Immunofluorescence | Anti-Tyrosinase (M) | 1/200 | Bio-techne NBP2-67233 |
| Immunofluorescence | Secondary Donkey anti-Rabbit IgG (H+L) Antibody, Alexa Fluor™ 488 | 1/500 | Invitrogen A21206 |

### Histology and immunofluorescence staining

Human and mice tissue samples were fixed in formalin for 24 h at room temperature and paraffin-embedded. Formalin-fixed paraffin-embedded (FFPE) tissues were sectioned at 4 µm using a microtome and mounted onto SuperFrost Plus adhesion slides. Sections were dried at approximately 56°C for at least 2 h and subsequently stabilized for at least 24 h at room temperature before staining.

For hematoxylin and eosin (H&E) staining, sections were deparaffinized in toluene, rehydrated through a graded ethanol series, and stained with hematoxylin and eosin according to standard procedures ^52,53^. For immunofluorescence, FFPE sections were heated at 60°C for 15 min, deparaffinized in xylene (3 × 5 min), and rehydrated through a graded ethanol series (100%, 95%, 70%, and 50% ethanol) before rising in distilled water. Heat-induced epitope retrieval (HIER) was performed using a PT-Link preprocessing system (Dako) with antigen-retrieval buffer at pH 6.0 or pH 8.0, depending on the primary antibody. Sections were heated from 70 °C to 97 °C over 20 min, maintained at 97 °C for 20 min, and cooled to 70 °C over 20 min. Following antigen retrieval, sections were washed twice in PBS for 5 min. Sections were permeabilized in PBS containing 0.01–0.05% Tween-20 for 5 min and blocked with 5% bovine serum albumin (BSA) in PBS for 1 h at room temperature. Sections were then incubated overnight at 4 °C with primary antibodies diluted in blocking solution. Antibody sources, catalog numbers, dilutions, and antigen-retrieval conditions are provided in **the above table**. After washing, sections were incubated with the appropriate fluorophore-conjugated secondary antibodies for 1 h at room temperature, counterstained with DAPI, and mounted using antifade mounting medium. Fluorescence images were acquired using an ECHO Revolution fluorescence microscope.

### Fontana-Masson Staining

Fontana–Masson staining was performed to visualize melanin deposition and distribution in skin sections. FFPE sections were deparaffinized and rehydrated as described above for H&E staining. Sections were incubated with preheated silver nitrate solution at 60 °C for 75 s. Slides were then rinsed three times in distilled water and toned with 0.2% gold chloride for 10 min at room temperature. Excess solution was removed, and sections were treated with 3% sodium thiosulfate for 2 min to remove unreacted silver. After two washes in distilled water, sections were dehydrated through graded ethanol, cleared in toluene, and mounted with Eukitt mounting medium.

Bright-field images were acquired at ×10 and ×40 magnification to evaluate melanin deposition and its spatial distribution within the epidermis and along the dermo–epidermal junction.

### Multiplex quantification of inflammatory mediators

Cytokine and chemokine concentrations in cell culture supernatants were quantified using the LEGENDplex™ Human Inflammation Panel (BioLegend, San Diego, CA, USA), which simultaneously measures IL-1β, IFN-α2, IFN-γ, TNF, MCP-1 (CCL2), IL-6, IL-8 (CXCL8), IL-10, IL-12p70, IL-17A, IL-18, IL-23, and IL-33. Assays were performed according to the manufacturer’s instructions. Samples were acquired on a FACSCanto™ II flow cytometer (BD Biosciences, Franklin Lakes, NJ, USA), and cytokine and chemokine concentrations were determined using the QOGNIT LEGENDplex™ data analysis software, as previously described ^18^.

### Chemical quantification of eumelanin and pheomelanin by HPLC-DAD

Melanin degradation products were prepared from mouse ear skin samples (n = 7 per group) by alkaline hydrogen peroxide oxidation. The surface area of each sample was measured before processing to enable normalization of melanin content to tissue area. Tissue samples were weighed, transferred to gentleMACS M tubes, and homogenized in 400 µL of 1 M KOH using a gentleMACS Dissociator (Miltenyi Biotec, Paris, France). Homogenates were centrifuged at 3,000 × *g* for 3 min and transferred to 15-mL screw-cap conical tubes. Melanin oxidation was performed by adding 25 µL of 30% (v/v) H_2_O_2_ and incubating overnight at 25 °C under continuous gentle agitation. At the end of the reaction, 50 µL of 10% (w/v) Na_2_SO_3_ was added, the pH was adjusted to 4.5–5.5 with 1 M HCl, and 675 µL of 100 mM sodium acetate buffer (pH 5.2) was subsequently added.

Melanin degradation products were purified by weak anion-exchange solid-phase extraction (SPE), as previously described ^54^. Strata-X-AW polymeric weak anion-exchange cartridges (33 µm, 30 mg/1 mL; Phenomenex) were sequentially conditioned with 500 µL of HPLC-grade acetonitrile and 500 µL of 100 mM sodium acetate buffer (pH 5.2). Tissue degradation extracts were loaded onto the conditioned cartridges at a reduced flow rate. Cartridges were washed sequentially with 500 µL of 100 mM sodium acetate buffer (pH 5.2) and 500 µL of acetonitrile and then dried under continuous vacuum for 5 min. Target analytes were eluted twice with 500 µL of 5% (v/v) aqueous NH_4_OH. The combined eluates were frozen and lyophilized overnight using an Alpha 2-4 LD Plus freeze dryer (Christ). Dried residues were reconstituted in 200 µL of 20 mM aqueous potassium phosphate buffer (pH 3.3).

Melanin degradation products were quantified using an Agilent 1200 HPLC system equipped with an online vacuum degasser, quaternary pump, autosampler, thermostated column compartment, and diode-array detector (DAD), controlled using OpenLab CDS software (v.A.01.05; Agilent Technologies). Chromatographic separation was performed on a Gemini C18 column (150 × 3 mm, 3 µm particle size, 100 Å pore size; Phenomenex) equipped with a SecurityGuard cartridge for Gemini C18. The mobile phase consisted of 20 mM aqueous potassium phosphate buffer (pH 3.3; solvent A) and HPLC-grade methanol (solvent B). Isocratic elution at 0% B was maintained from 0 to 6 min, followed by a linear increase from 0% to 10% B between 6 and 7 min. The column was subsequently re-equilibrated at 0% B from 7 to 15 min. The flow rate was maintained at 0.6 mL/min, the column temperature at 36 °C, and the injection volume was 20 µL.

The eumelanin degradation markers pyrrole-2,3,5-tricarboxylic acid (PTCA) and pyrrole-2,3-dicarboxylic acid (PDCA) were monitored at 225 nm and 280 nm, respectively. The pheomelanin degradation markers thiazole-2,4,5-tricarboxylic acid (TTCA) and thiazole-4,5-dicarboxylic acid (TDCA) were monitored at 275 nm and 254 nm, respectively. Peak identities were confirmed by comparison of retention times and UV-visible absorption spectra with those of purified reference standards. Eumelanin content was estimated from the quantified PTCA amount using a conversion factor of 60, whereas pheomelanin content was estimated from the quantified TTCA amount using a conversion factor of 34. Final eumelanin and pheomelanin contents were normalized to the surface area of the corresponding tissue sample and expressed as ng/cm^2^.

### Transmission Electron Microscopy (TEM)

Mouse ear tissue fragments were collected from the respective experimental groups (n = 3 mice per group) and initially fixed in 2% (v/v) paraformaldehyde and 2.5% (v/v) glutaraldehyde in 0.1 M Sorensen buffer (pH 7.4) for 2 h at room temperature, followed by further fixation for 1 week at 4 °C. Samples were subsequently fixed in fresh 2.5% (v/v) glutaraldehyde in 0.1 M Sorensen buffer (pH 7.4) using a PELCO BioWave Pro+ microwave system (Ted Pella, Inc.). Samples were immersed in fixative for at least 15 min and subjected to microwave irradiation under vacuum at 150 W using a 2-min on, 2-min off, and 2-min on cycle. After rinsing in 0.1 M Sorensen buffer, samples were post-fixed in 1% osmium tetroxide (OsO_4_) using microwave-assisted processing under vacuum at 150 W for 30 min, with alternating 3-min irradiation and 2-min pause cycles. Samples were then dehydrated using a graded acetone series (50%, 70%, 95%, 95%, 100%, and 100%) in the microwave at 200 W for 1 min per step. Resin infiltration was performed using graded mixtures of Embed 812 resin and acetone (1:3, 1:1, and 3:1) under vacuum at 300 W for 4 min per step, followed by three successive infiltrations in 100% resin for 5 min each at 300 W. Samples were embedded in fresh resin and polymerized at 60 °C for 48 h.

Ultrathin sections (70 nm) were cut using a diamond knife (Diatome) on an EM UC7 ultramicrotome (Leica Microsystems), collected onto Butvar-coated copper grids, and post-stained with UranyLess and lead citrate. Sections were examined using an H7650 transmission electron microscope (Hitachi) operated at 80 kV in high-contrast mode. Images were acquired using a 4K × 4K RIO16 CMOS camera (Gatan) and processed using DigitalMicrograph software (Gatan). Epidermal and dermal compartments were systematically examined across multiple tissue sections. Melanocytes identified within the sections were subsequently imaged at higher magnification for quantitative assessment of melanosome content. To ensure balanced sampling across experimental groups, 55 melanocytes per condition were randomly selected and analyzed. Statistical comparisons were performed using Welch’s ANOVA followed by the Games–Howell multiple-comparison test.

### Xenium 5K spatial transcriptomic analysis

#### Tissue processing and data acquisition

Ear skin samples from the four experimental groups were processed as formalin-fixed paraffin-embedded (FFPE) tissues according to the 10x Genomics Xenium FFPE tissue preparation workflow. Tissue sections (5 µm) were analyzed in two independent experimental batches performed three months apart. Each batch comprised two Xenium slides: one slide containing eight irradiated samples (four *Xpc*^+/+^ IR and four *Xpc*^−/−^ IR) and a second slide containing eight non-irradiated samples (four *Xpc*^+/+^ NIR and four *Xpc*^−/−^ NIR). Thus, Xenium spatial transcriptomic profiling was initially performed on eight mouse ear samples per experimental condition. Following quality control, 27 samples were retained for downstream analyses for the non-irradiated *Xpc*^+/+^ (n=6), irradiated *Xpc*^+/+^ (n=8), non-irradiated *Xpc*^−/−^ (n=6), and irradiated *Xpc*^−/−^ (n=7) groups.

Gene detection was performed using the pre-designed 10x Genomics Xenium 5K Mouse Pan Tissue panel supplemented with a custom 100-gene add-on panel. The custom panel comprised 50 genes selected to improve cell-type and subpopulation resolution, particularly within the keratinocyte and immune-cell compartments, and 50 genes associated with pigmentation and metabolic pathways for downstream signature analyses. FFPE tissue sections (5 µm) were placed on Xenium slides and processed on the Xenium Analyzer according to the 10x Genomics Xenium In Situ Gene Expression protocols (CG000580_RevE and CG000760_RevC). The Xenium 5K Mouse Pan Tissue and custom panels were hybridized to the tissue sections for 18 h overnight. Tissue sections were subsequently stained according to the Xenium Cell Segmentation workflow, and slides were loaded onto the Xenium Analyzer for imaging and decoding of spatially resolved transcripts.

Cell segmentation was performed using the multimodal 10x Genomics Xenium segmentation algorithm. This deep-learning-based approach integrates DAPI nuclear staining with E-cadherin/ATP1A1/CD45, and αSMA/vimentin fluorescence signals to delineate cellular boundaries. Spatial visualization and inspection of tissue and cell distributions were performed using Xenium Explorer (10x Genomics).

#### Data processing and cell-type annotation

Following spatial decoding, Xenium output data, including transcript coordinates, quality scores, and cell boundaries, were assembled into AnnData objects and analyzed using Scanpy in Python. Cells containing fewer than 80 total transcripts or fewer than 10 detected genes were excluded. Transcript counts were normalized to 10,000 counts per cell and log-transformed as ln(counts + 1).

Highly variable genes were selected, and principal component analysis (PCA) was performed on the scaled expression matrix. Technical variation between slides and experimental batches was corrected using Harmony. Harmony-corrected principal components were used to construct a nearest-neighbor graph, followed by graph-based clustering using the Leiden algorithm.

Clustering was performed across a range of resolutions (0.1–1.9, in increments of 0.2), and the final resolution was selected based on cluster stability, separation, and biological interpretability. Cluster stability was evaluated using clustree ^55^, which visualizes the relationships between clusters across successive resolutions and tracks changes in cell assignments. The lowest resolution at which a distinct cluster composed predominantly of melanocytes, with contributions from all experimental samples, could be identified was selected for subsequent analyses (resolution = 1.1). The selection of resolution 1.1 was further supported by the mean silhouette score, which provided an independent assessment of cluster separation, with higher values indicating better-defined clusters. Biological interpretability was assessed by examining the top differentially expressed genes for each cluster to determine whether they supported the identification of distinct cell populations. Finally, clusters were manually annotated based on established cell-type-specific marker genes and their spatial distribution within skin tissue. Marker selection and cell-type annotation were guided by published skin single-cell RNA-sequencing reference datasets available through the Human Protein Atlas ^56^.

#### Spatial and differential expression analyses

Ear sections were divided into basal, middle, and distal regions to enable spatially resolved comparisons across experimental groups. Regional boundaries were defined based on anatomical position and tissue morphology (i.e. epidermal thickness) and were subsequently applied consistently across samples. For each sample and anatomical region, the melanocyte-to-keratinocyte ratio was calculated as the number of annotated melanocytes divided by the number of annotated keratinocytes, and statistical comparisons were performed using pairwise Mann–Whitney U test.

Differential gene expression analysis was performed using the Wilcoxon rank-sum test with Benjamini–Hochberg correction for multiple testing. Genes with an adjusted *P* value (FDR) < 0.05 and an absolute log_2_ fold change > 1 were considered significantly differentially expressed. Functional and pathway enrichment analyses were performed using IPA (QIAGEN). IPA was additionally used to identify canonical pathways and predict their activation states and associated biological functions based on activation z-scores.

### Hyperion imaging mass cytometry

#### Antibody metal conjugation

Antibodies were conjugated to metal isotopes using Maxpar labeling kits according to the manufacturer’s instructions. The antibody labeled with a cadmium isotope was conjugated using the Maxpar® MPC9 Antibody Labeling Kit, whereas all other antibodies were labeled using the Maxpar® X8 Antibody Labeling Kit (PRD002 Rev 14; Fluidigm/Standard BioTools). Details of the antibodies and corresponding metal isotopes are provided in **supplementary Table S2**.

#### Tissue labelling

FFPE tissue sections (5 µm) were deparaffinized and subjected to heat-induced antigen retrieval using Dako Target Retrieval Solution, pH 9 (S236784-2; Agilent Technologies), in a water bath at 96 °C for 30 min. Tissue sections were encircled using a Dako Pen and incubated with SuperBlock™ Blocking Buffer (37515; Thermo Fisher Scientific) for 45 min at room temperature, followed by Fc receptor blocking reagent (130-092-575; Miltenyi Biotec) for 1 h at room temperature.

For tyrosinase detection, sections were incubated with the primary anti-tyrosinase antibody overnight at 4 °C. After washing, tyrosinase was detected using a ^141^Pr-conjugated anti-mouse secondary antibody for 2 h at room temperature. Sections were then washed three times for 8 min each in PBS containing 0.2% Triton X-100 (PBS-T).

The metal-conjugated antibody cocktail (**Table S2**) was diluted in PBS containing 1% BSA and applied to the tissue sections overnight at 4 °C. Sections were subsequently washed three times for 8 min each in PBS-T. Nuclei were labeled with an iridium-based DNA intercalator (1:400 in PBS; Fluidigm/Standard BioTools) for 30 min at room temperature. Finally, sections were washed for 5 min in PBS followed by 5 min in distilled water and air-dried at room temperature for 30 min before imaging.

#### Imaging mass cytometry: data acquisition

Images were acquired using the Hyperion™ XTi Imaging System (Fluidigm/Standard BioTools) according to the manufacturer’s instructions. After selection of regions of interest (ROIs), tissue sections were ablated using a UV laser at a frequency of 800 Hz. Acquired data were exported as MCD files and visualized using MCD™ Viewer software (v1.0.560.6; Fluidigm/Standard BioTools). For visualization, minimum and maximum display thresholds were adjusted for each marker and tissue section to optimize signal visualization, while the gamma value was maintained at 1.

### General statistical analysis

Unless otherwise specified in the respective Methods sections, statistical analyses were performed using GraphPad Prism (v10.0; GraphPad Software) and Perseus (v2.1.6.0). Pairwise comparisons between two groups were performed using two-tailed Student’s *t*-test or the Mann–Whitney U test, as appropriate. Comparisons involving multiple groups were performed using two-way analysis of variance (ANOVA), as appropriate, followed by Šídák’s multiple-comparison test. Data are presented as mean ± SD unless otherwise indicated. Statistical significance was defined as *P* < 0.05 and represented as follows: \**P* < 0.05, \*\**P* < 0.01, \*\*\**P* < 0.001, and \*\*\*\**P* < 0.0001; ns, not significant (*P* ≥ 0.05). Exact *P* values are reported where applicable. The number of independent biological replicates (*n*) and the statistical test used for each experiment are specified in the corresponding figure legends. Statistical procedures specific to proteomic, metabolomic, kinome, and spatial transcriptomic analyses, including multiple-testing corrections, are described in the respective Methods sections.

